# Dlg5 and Cadherins are key to peripheral glia integrity

**DOI:** 10.1101/2022.11.01.514384

**Authors:** Das Mriga, Cheng Duo, Matzat Till, Auld Vanessa J.

## Abstract

Glial cells in the peripheral nerve wrap axons to insulate them and ensure efficient conduction of neuronal signals. In the myelin sheath, it is proposed that the autotypic tight junctions and adherens junctions form glia-glia complexes that stabilize the glia sheath in myelinating glia. Yet the role of adhesion junctions in non-myelinating glia of vertebrates or invertebrates has not been clearly established. Many components of adhering junctions contain PDZ (PSD-95, Dlg, ZO1) domains or are recruited to these junctions by PDZ binding motifs. To test for the role of PDZ domain proteins in glial sheath formation, we carried out an RNAi screen using *Drosophila melanogaster* to knockdown each of the 66 predicted PDZ domain proteins in the peripheral glia. We identified six PDZ genes with potential roles in glial morphology, and further investigated Discs-large 5 (Dlg5), a scaffolding protein with no previously known function in glia. Knockdown of Dlg5 disrupts subperineurial glia (SPG) morphology, including gaps in the membrane that coincide with disruption of septate junction proteins. To further our investigation of Dlg5, we focused on cadherins and found both N-Cadherin and E-Cadherin are expressed throughout peripheral glia. Knockdown of E-Cadherin phenocopied the loss of Dlg5 leading to gaps in the SPG and septate junctions while only simultaneous loss of both N-Cadherins (NCad, and CadN2) had the same effect. The loss of all three Cadherins enhanced these phenotypes as did loss of Dlg5 when paired with cadherin knockdown. This leads to a model where Dlg5 plays a role in conjunction with cadherins in glial membrane stabilization and septate junction formation in the subperineurial glia.

## INTRODUCTION

A critical function of glia is to wrap and ensheath axons in both the central and peripheral nervous systems. In vertebrate peripheral nerves, myelinating Schwann cells enable rapid and efficient transduction of electrical signals by forming the myelin sheath to ensheath and insulate large caliber axons. While there are extensive studies that have identified the protein complexes that mediate myelin sheath formation and stabilization, almost nothing is known about the adherent junctions that mediate sheath formation in non-myelinating Schwann cells. Yet this class of glia is essential to protect and support small calibre axons such as C-fibers that transmit nociceptive information (Chen et al., 2003; Griffin and Thompson, 2008).

Many components of adherent junctions such as tight junctions (TJ) are either PDZ (PSD-95, Dlg, ZO1) proteins or contain motifs allowing for interactions with PDZ proteins. PDZ domains are 80-90 amino acids and mediate protein-protein interaction by binding to short three amino acid PDZ binding motifs found at the C-terminus of many proteins (Harris and Lim, 2001; Subbaiah et al., 2011). Many PDZ proteins possess different combinations of protein interaction domains that mediate the recruitment of protein complexes. For instance, the family of membrane-associated guanylate kinases (MAGUKs) apart from PDZ domains, possess L27, SH3 and GUK domains. PDZ proteins regulate processes such as cell polarity, adhesion, migration and signalling through their ability to scaffold proteins in specific regions of the cell. It is likely that PDZ proteins play a role in mediating non-myelinating glia junctions and are critical for glial cell function.

Acquiring a better understanding of PDZ proteins and their role in glial junctions requires that we identify those PDZ proteins that function in non-myelinating glia. Our model is the peripheral nerve of *Drosophila melanogaster* given that there are only 66 PDZ proteins in Drosophila (Aranjuez et al., 2012 2001) (compared to the 250 predicted PDZ proteins in humans (Tonikian et al., 2008)) and that *Drosophila* peripheral glia are an excellent model to further our understanding of the mechanisms underlying non-myelinating glia development and function. Each peripheral nerve is ensheathed by three glial layers. The innermost glia layer, the wrapping glia (WG) resemble non-myelinating Schwann cells and extend their processes around individual or bundles of motor and sensory axons. The wrapping glia layer is surrounded by the subperineurial glia (SPG), which form septate junctions between the glial layers to establish the blood-nerve barrier to protect peripheral axons from the high K+ hemolymph (Auld et al., 1995; Baumgartner et al., 1996). Septate junctions form a ladder-like array of a core protein complex that is conserved with paranodal junctions including Neurexin IV, Neuroglian and Contactin (Bhat, 2003; Limmer et al., 2014). Lastly, the perineurial glia (PG), which are the equivalent to vertebrate perineurial cells, wrap the nerve and adhere to the extracellular matrix that surrounds and stabilizes the entire nerve. Adhesion and signalling between these different glial layers must require glia-glia interactions, however the proteins and in particular the PDZ proteins required for these processes in remain largely unknown.

In this study, we screened all 66 Drosophila PDZ-encoding proteins using RNAi knockdown to determine which play a role in peripheral glia. We identified seven PDZ domain proteins where loss of function lead to disruption of peripheral glia and focused our subsequent investigation on Discs-large 5 (Dlg5). Dlg5 is a MAGUK protein with a coiled-coil domain at the amino terminus, four PDZ domains, an SH3 and GUK domain, and has been implicated in cell polarity and Cadherin complex formation (Nechiporuk et al., 2007; Wang et al., 2014; Reilly et al., 2015; Liu et al., 2017). The role of Dlg5 and those of the classical Cadherins in non-myelinating glia, including the peripheral glia of Drosophila, had not been determined. Therefore, we went onto investigate Dlg5 and potential functional overlap with the Drosophila classical cadherins; E-Cadherin (ECad, *shotgun*), N-Cadherin (NCad, *CadN*), N-Cadherin2 (NCad2, *CadN2*) in the three layers of peripheral glia. Knockdown of Dlg5 in the subperineurial glia resulted in breaks/gaps within the glial membrane and disruption of septate junction proteins. We found that ECad, NCad and CadN2 perform redundant functions in the subperineurial glia, and that loss of Cadherins phenocopied and enhanced the loss of Dlg5 phenotypes. Overall, our results show that Dlg5 interacts with cadherins to mediate SPG and septate junction morphology.

## MATERIALS AND METHODS

### Fly strains and genetics

The following fly strains were used in this study: repo-GAL4 (Sepp et al., 2001); Nrv2-GAL4 (Sun et al., 1999), 46F-GAL4 (Xie and Auld, 2011); Gli-GAL4 (Sepp and Auld, 1999); SPG-GAL4 (Schwabe et al., 2005); UAS-mCD8::GFP (Lee and Luo, 1999); UAS-Dicer2 (Dietzl et al., 2007); *Df(2L)BSC148*, UAS-mCD8::RFP, UAS-Dlg5::GFP (Bloomington Stock Center, BDSC). The following GFP and RFP protein-trap insertions were used: Dlg5::GFP (Sarov et al., 2016); Nrv2::GFP, NrxIV::GFP, Scrib::GFP (Morin et al., 2001; Kelso et al., 2004; Buszczak et al., 2007); NCad::cherry, ECad::cherry (Venken et al., 2011). The following RNAi lines were used and those lines preferentially used for the majority of experiments are in bold: Dlg5-RNAi (VDRC: **46234**, 101596; BDSC: 41832); Arm-RNAi (VDRC: 7767, **107344**); ECad-RNAi (VDRC: **27802**; 103962; BDSC: 27689, 32904, 38207); NCad-RNAi (VDRC: **1092**; BDSC: 27503); CadN2-RNAi (VDRC: 101659; BDSC: **27508**); Crumbs-RNAi (VDRC: **39177**). All RNAi experiments carried out after the initial screen were raised at 25°C either with or without Dicer2 in the background. For all experiments, controls were the driver line crossed wit *w[1118]*.

#### Screen of RNAi knockdown of PDZ genes

RNAi lines were obtained from TRiP and VDRC stock centers. Table 1 provides the list of RNAi lines used in the screen. *repo>mCD8::RFP/TM6,Tb* virgin flies were crossed with males from each UAS-RNAi line at 29°C. As a control, *repo>mCD8::RFP/TM6,Tb* virgin flies were crossed with *w^1118^* males and raised at 29°C. A minimum of 6 larvae were dissected from each cross and stained with mAb 22C10 (anti-Futsch) to label the peripheral axons. The morphology of the glial membranes were visualized using UAS-mCD8::RFP.

#### Immunolabeling and image analysis

The peripheral nervous system of 3^rd^ instar, wandering larvae were dissected and fixed for immunolabeling using previously described methods (Sepp et al., 2000). The following primary antibodies were used in this study: guinea pig anti-Inx2 (1:500) (Smendziuk et al., 2015); mouse anti-Futsch/22C10 (1:1000, DSHB); rabbit anti-GFP (1:600, Life Technology); mouse anti-GFP (1:300, Novus Biologicals), rabbit anti-Cherry (1:300, Abcam), mouse anti-βPS (1:50, CF.6G11) (Brower, 1984), rabbit anti-HRP (1:500, Jackson ImmunoResearch, West Grove, PA); mouse anti-Futsch/22C10 (1:1000, DSHB); rabbit anti-Nrv2.1 (1:1000, Abcam). The following secondary antibodies were used at a 1:300 dilution: goat anti-mouse Alexa 488, Alexa 568 and Alexa 647; goat anti-rabbit Alexa 568 and goat anti-guinea pig Alexa 647 (Molecular Probes). DAPI (1:1000, Invitrogen) was used to label nuclei.

Images were acquired with a Delta Vision Spectris compound microscope (Applied Precision/GE Healthcare, Mississauga, Ontario) using a 60x oil immersion objective (NA 1.4). An image was captured every 0.2 μm and the resulting stacks were deconvolved (SoftWorx, Toronto, Canada) using a point spread function measured with 0.2 μm beads conjugated to Alexa dyes (Molecular Probes) and mounted in Vectashield (Vector Laboratories, Burlington, Canada). Cross sections were generated using SoftWorx. A single z-slice was chosen from each z-stack and images were compiled using Adobe Photoshop and Adobe Illustrator CC. For transmission electron microscopy analysis larvae were dissected and prepared using previously described methods (Matzat et al., 2015) using a Hitachi H7600 TEM at the UBC BioImaging Facility.

#### Dye Penetration Assay

Third instar larvae were washed using 1X PBS, then opened at the posterior end and inverted in Schneider’s insect medium (Millipore Sigma, S0146). The inverted larvae were incubated in Schneider’s medium containing 2.5 mM of 10,000MW Dextran Texas Red-conjugated dye (Invitrogen, D1863, Lysine Fixable) for 40 minutes at room temperature on an orbital shaker. The larvae were then fixed using 4% PFA with 1X PBS for 20 minutes. Following fixation, samples were washed with 1X PBS twice for 5 minutes each, then washed with 0.1% PBST with 1X PBS twice for 10 minutes each. Lastly, the brains were dissected and mounted in Vectashield. The samples were imaged using an Olympus FV1000 Confocal microscope (BioImaging Facility UBC) using a 30X Silicone oil immersion objective (NA = 1.05) with 1um steps. Laser (559nm) intensity was set at 7% intensity, 750V digital gain, 1x master Gain, and 6% offset for all genotypes except for imaging of NrxIV-RNAi, which was 5% intensity, 750V digital gain, 1x master Gain, and 6% offset. Images were imported into ImageJ and fluorescence intensity calculated and images displayed using the Fire Look up Table.

#### Locomotion assay

Locomotion assays were performed using third instar larvae adapted from (Brooks et al., 2016). Larvae of the desired genotype were gently placed onto a freshly made 2% agar plate with food-safe dye added to enhance contrast. Larval movements were recorded continuously for 60 seconds using a Canon VIXIA HF R800 video camera. The recordings were then converted from *.MOV to *.AVI files using FFmpeg (ffmpeg.org) and analyzed using the wrMTrck plug-in (Nussbaum-Krammer et al., 2015; Brooks et al., 2016) in Fiji to calculate locomotion and movement trajectory. Larvae with non-continuous paths throughout the entirety of the recording were discarded from the total travel distance quantification but were included in the average travel speed.

#### Statistics

Prism (GraphPad Software) was used for all statistical analyses and a One-way Anova utilized to compared multiple genotypes with the specified control with a Dunnet multiple comparison test. For comparisons across multiple genotypes a Tukey multiple comparison test was used. For TEM data analysis a Kruskal-Wallis non-parametric test with a Dunn’s multiple comparisons test was utilized. For all fluorescence image analyses at least 6-7 nerves were analyzed per larvae.

## RESULTS

### RNAi induced knockdown of PDZ proteins in glial cells

To understand the role of PDZ proteins in mediating glia interactions, we utilized the peripheral nerve of the 3^rd^ instar *Drosophila* larva to investigate the consequences on glial morphology of knockdown of PDZ domain proteins. There are 66 genes predicted to encode PDZ domain proteins in the *Drosophila* genome (Aranjuez et al., 2012). Using the pan-glial driver *repo-GAL4*, we expressed each *UAS-PDZ-RNAi* line while co-expressing *UAS-mCD8::RFP* to label the glial membranes. We scored morphological defects in the labeled peripheral glial membranes and axons were labeled using anti-Futsch to reveal any axonal defects. When available, multiple RNAi lines were used to target each PDZ gene. For the 66 genes, 150 individual *UAS-RNAi* lines were tested for abnormal glial phenotypes in the PNS (Fig.1A, Supplemental Table 1). As a control *repo>mCD8::RFP* flies were crossed to *w^1118^* flies (Fig. 1B-B’). We classified PDZ genes as positive hits (Fig. 1A; Supplemental Table 1) if their knockdown using multiple RNAi lines resulted in lethality or glial phenotypes. Seven PDZ proteins; Discs Large 1 (Dlg1), Scribbled (Scrib), Varicose (Vari), Stardust (sdt), Discs Large 5 homolog (Dlg5), Locomotion defects (loco), and a Rho-type guanine nucleotide exchange factor (RhoGEF2) were identified as positive hits with potential functions in glial cells (Fig. 1A,C-G; Supplemental Table 1). The screen resulted in a range of phenotypes such as glial swelling observed with the knockdown of RhoGEF2 and Loco (Fig. 1C-C”, F-F”). Knockdown of Dlg1 and Scrib resulted in abnormal axonal wrapping (Fig.1 D-D”, E-E”), whereas Vari knockdown in all glia was lethal (Table 1). Abnormalities in axonal wrapping, specifically by the inner wrapping glial membrane were observed with Dlg5 knockdown (Fig. 1G-G’). Of note is that loss of known apical polarity proteins including Bazooka (baz, Par-3), Par-6 and Patj had no effect (Fig. 1A; Supplemental Table 1). Conversely the basolateral polarity proteins such as Dlg1 and Scrib had strong effects as did Dlg5, which is also thought to function in epithelial polarity (Luo et al., 2016). Overall, we identified a group of PDZ proteins that may play necessary roles in peripheral glia development and a number of these proteins are known to function in polarity and junction formation.

**Figure 1:**
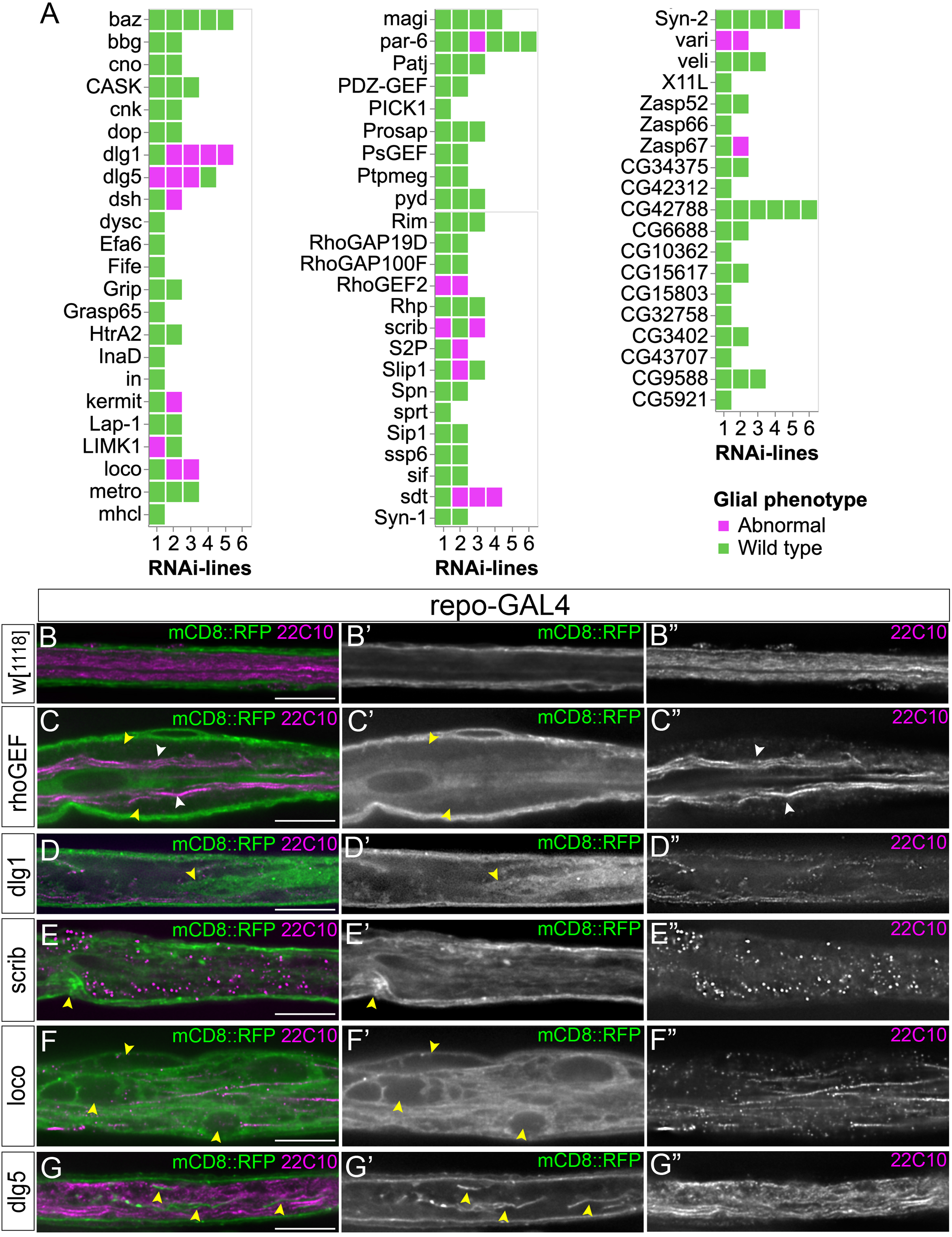
Identification of PDZ proteins required in glial cells of the peripheral nerve. **A:** Summary of PDZ protein RNAi screen. Details of RNAi lines used to target each candidate gene can be found in Table 1. Each box represents an individual RNAi line with green meaning wild type and magenta representing abnormal glial phenotypes. **B-G:** Longitudinal sections of peripheral nerves from control and selections of positive candidate identified in the RNAi screen. All glial membranes labeled with mCD8::RFP (green) and axons immunolabeled with anti-Futsch (22C10; magenta). The pan-glial driver repo-GAL4 was used and crossed to: (B-B”) Control (*w[1118]*) peripheral nerve. (C-C”) Rho-GEF2-RNAi. Swollen regions are indicated (C’, yellow arrowheads) and these swellings were accompanied by defasciculation of the axons in that region (white arrowheads, C’’). (D-D”) Dlg1-RNAi. Glial membranes were disrupted (D’, yellow arrowhead). (E-E”) Scrib-RNAi. Glial membranes were abnormal with accumulations (E’, yellow arrowhead). (F-F”) Loco-RNAi. Peripheral nerves were swollen and vacuole-like structures were observed within the glial membranes (F’, yellow arrowheads). (G-G”) Dlg5-RNAi. Peripheral nerves had disrupted inner glial membranes (G’, yellow arrowheads). The morphology of axons in these nerves were unaffected (G’’). Scale bars: 15μm

### Dlg5 is expressed in glial cells of the larval peripheral nerve

Given the role of Dlg5 in cell polarity and cell-cell adhesion, and the fact that Dlg5 has not been studied in glia, we chose to investigate Dlg5 further. Dlg5 is a MAGUK protein with a coiled-coil domain at the amino terminus, four PDZ domains, an SH3 and GUK domains. We began by examining the distribution of Dlg5 in the glial layers of the peripheral nerve in 3rd instar larvae using Dlg5 endogenously tagged with GFP (Dlg5::GFP)(Sarov et al., 2016). Dlg5 puncta were observed throughout the peripheral nerve, with puncta present within the glial cell membranes (identified with mCD8::RFP) (Fig. 2A-A”). Some Dlg5 puncta might also be present in the axons (immunolabeled with anti-futsch) (Fig. 2B-B”). However, knockdown of Dlg5 using the pan-neuronal driver elav-GAL4, did not affect either peripheral glial or axon morphology (n=6 larvae).

**Figure 2.**
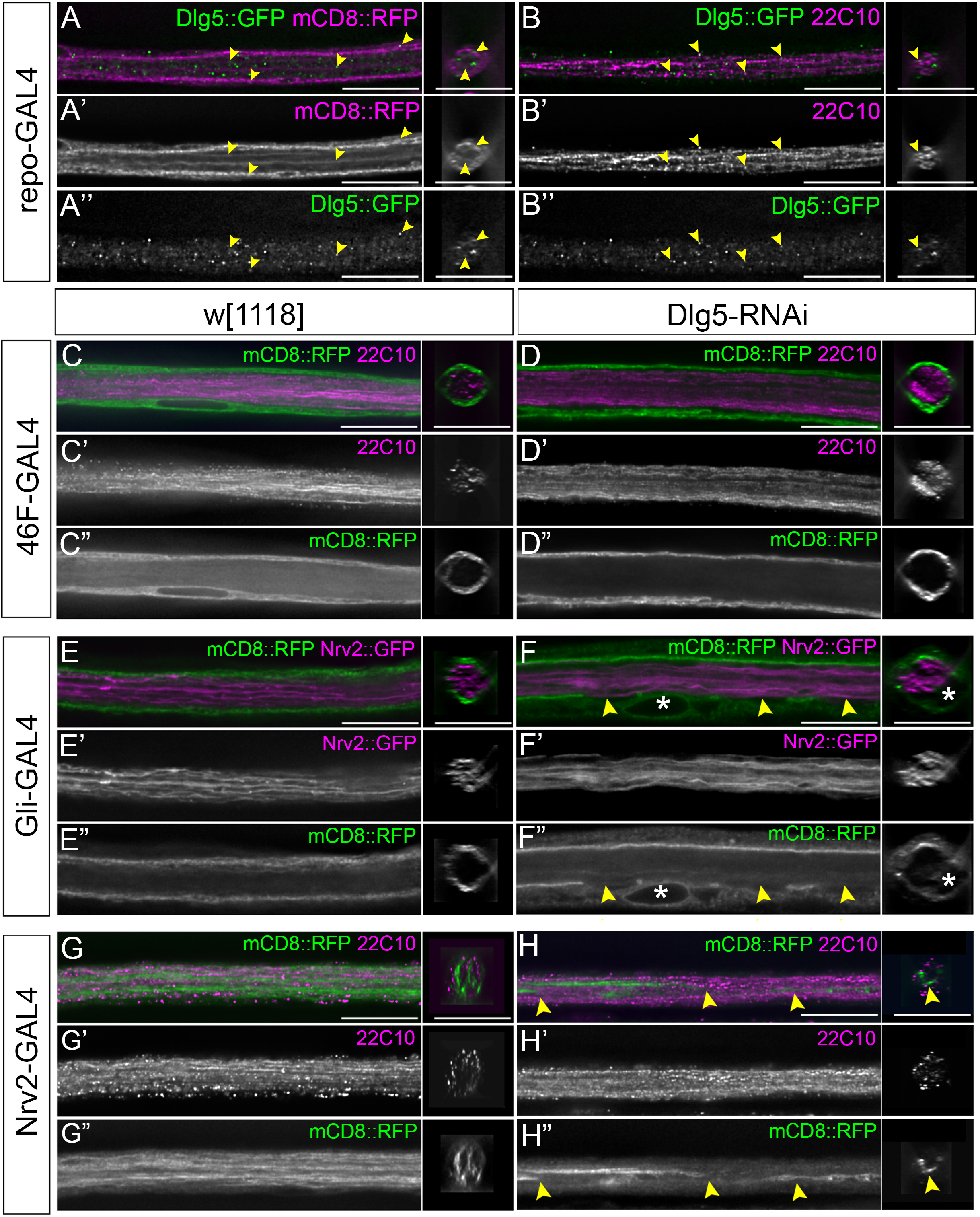
Knockdown of Dlg5 in the peripheral glial leads to disruptions of the SPG and WG. **A-B:** Endogenously tagged Dlg5 (Dlg5::GFP, green) reveals expression within glia and axons in the PNS. All glial membranes labeled with mCD8::RFP (repo-GAL4, magenta)(A) or with axons immunolabeled with anti-Futsch (22C10, magenta)(B). **C-D:** Perineurial glia. The PG driver 46F-GAL4 crossed to w[1118] (C) or Dlg5-RNAi (D). The PG membranes (mCD8::RFP, green) and axons (magenta) in both appeared normal in longitudinal and cross sections. **E-F**: Subperineurial glia. The SPG driver Gli-GAL4 crossed to w[1118](E) or Dlg5-RNAi (F) SPG membranes were labeled with mCD8::RFP (green) and WG membranes with Nrv2::GFP (magenta). The SPG membrane in control (**E’**) was continuous along the length of the nerve. In Dlg5-RNAi nerves (**G’**), the SPG was disrupted with breaks/gaps in the membrane (yellow arrowheads). The WG in both control (E’’) and Dlg5-RNAi (G’’) nerves extended normal processes along the length of the nerve. An SPG nuclei (asterisk) is indicated. **G-H**: Wrapping glia. The WG driver Nrv2-GAL4 crossed to w[1118] (G) and Dlg5-RNAi (**H**). The WG membranes were labeled with mCD8::RFP (green) and axons labeled with anti-Futsch (22C10, magenta). WG in control nerves (G”) extended normal processes along the length of the nerve and in cross-sections surrounded the axons (magenta). WG in Dlg5-RNAi nerves had fewer and disrupted processes (H”, yellow arrowheads). In cross-sections the WG processes did not extend around the axons (yellow arrowhead). Scale bars: 15μm

To further characterize the effects of Dlg5 loss in the peripheral glia, we knocked down Dlg5 in individual glial layers using GAL4 drivers specific to each of the three glial layers driving a membrane marker, mCD8::RFP. We tested multiple RNAi lines and used the strongest RNAi line, 46234, for this and future experiments. Knockdown of Dlg5 in the PG using 46F-GAL4 (*46F>mCD8::RFP,Dlg5-RNAi*) (n=5 larvae) did not affect PG membrane morphology (Fig. 2D-D”) compared to control (*46F>mCD8::RFP*)(Fig 2A-A”)(n=7 larvae). Knockdown of Dlg5 using Gli-GAL4 in the intermediate SPG layer (*Gli>mCD8::RFP,Dlg5-RNAi*) disrupted SPG morphology (Fig. 2F,F”), but had no effect on the WG (marked with Nrv2::GFP) (Fig. 2F’)(n=6 larvae). SPG membrane morphology was disrupted in 16.5% of nerves, with breaks/gaps in the membrane (yellow arrows) and occasionally accompanied by swellings (Fig. 2F,F”, asterisks). The control (*Gli>mCD8::RFP*) did not have any defects (Fig. 2E-E”)(n= 6 larvae). Knockdown of Dlg5 in the WG using Nrv2-GAL4 (*Nrv2>mCD8::RFP,Dlg5-RNAi*) resulted in reduced and discontinuous WG strands in 53% of nerves (Fig. 2H”) (n=5 larvae) compared to the control nerves (*Nrv2>mCD8::RFP*), where multiple, continuous WG strands ensheath the axons (Fig. 2G”)(n=6 larvae). Dlg5 knockdown in the WG did not affect axonal morphology (Fig. 2H’) which was similar to control nerves (Fig. 2G’). Overall, our results suggest the Dlg5 is expressed in multiple peripheral glia and is required in both the SPG and the WG but not in the PG layer.

### Dlg5 is required for the formation of septate junctions

Since Dlg5 knockdown affected SPG morphology, we next tested if the defects in the SPG extended to the septate junction (SJ) protein complex, which in Drosophila forms the blood-nerve and blood-brain barriers. Using a range of SJ proteins we looked for changes in the SJ morphology while knocking down Dlg5 in the SPG (*Gli>mCD8::RFP, Dlg5-RNAi*). For multiple RNAi lines (101596, 41832, 46234), we observed nerves with gaps in the SJ strands when labeled with NrxIV::GFP (Fig. 3B-B”) or immunolabeled with Nrv2.1 (Fig. 3D,D”) and often these corresponded to gaps in the mCD8::RFP labeled membrane (Fig. 3B”,D”) (n=12 larvae). In comparison, intact and continuous SJs were observed along the length of control nerves (*Gli>mCD8::RFP*)(Fig. 3A-A”,C-C”) (n=12 larvae). Increasing the efficiency of the RNAi with Dicer2 paired with a stronger SPG driver, moody-GAL4, increased the severity of the phenotypes (*SPG>mCD8::RFP,Dlg5-RNAi,Dicer2*) and NrxIV::GFP was often observed in large aggregates that corresponded to glial membrane concentrations (Fig. 3E-E”). We quantified the percentage of nerves per larvae (using moody-GAL4) that had disrupted SPG or gaps. On average 80% of the nerves were affected and with Dicer2, the penetrance was increased to 100% (Fig. 3F). To determine if the effect to Dlg5 in the SPG was cell autonomous, we knocked down Dlg5 in the PG and WG and assayed SJs using either *NrxIV::GFP* or the Nrv2.1 antibody. The morphology of SJs was not affected by the knockdown of Dlg5 in the PG layer (*46F>mCD8::RFP, Dlg5-RNAi*) (n=6 larvae). Similarly, knockdown of Dlg5 in the WG (*NrxIV::GFP, Nrv2>mCD8::RFP, Dlg5-RNAi*) lead to the expected WG phenotypes, but did not affect the morphology of the SJ strands (n= 6 larvae). Therefore, loss of Dlg5 in the SPG leads to a range of defects in the SPG including disruption of the SJs in a cell autonomous manner.

**Figure 3.**
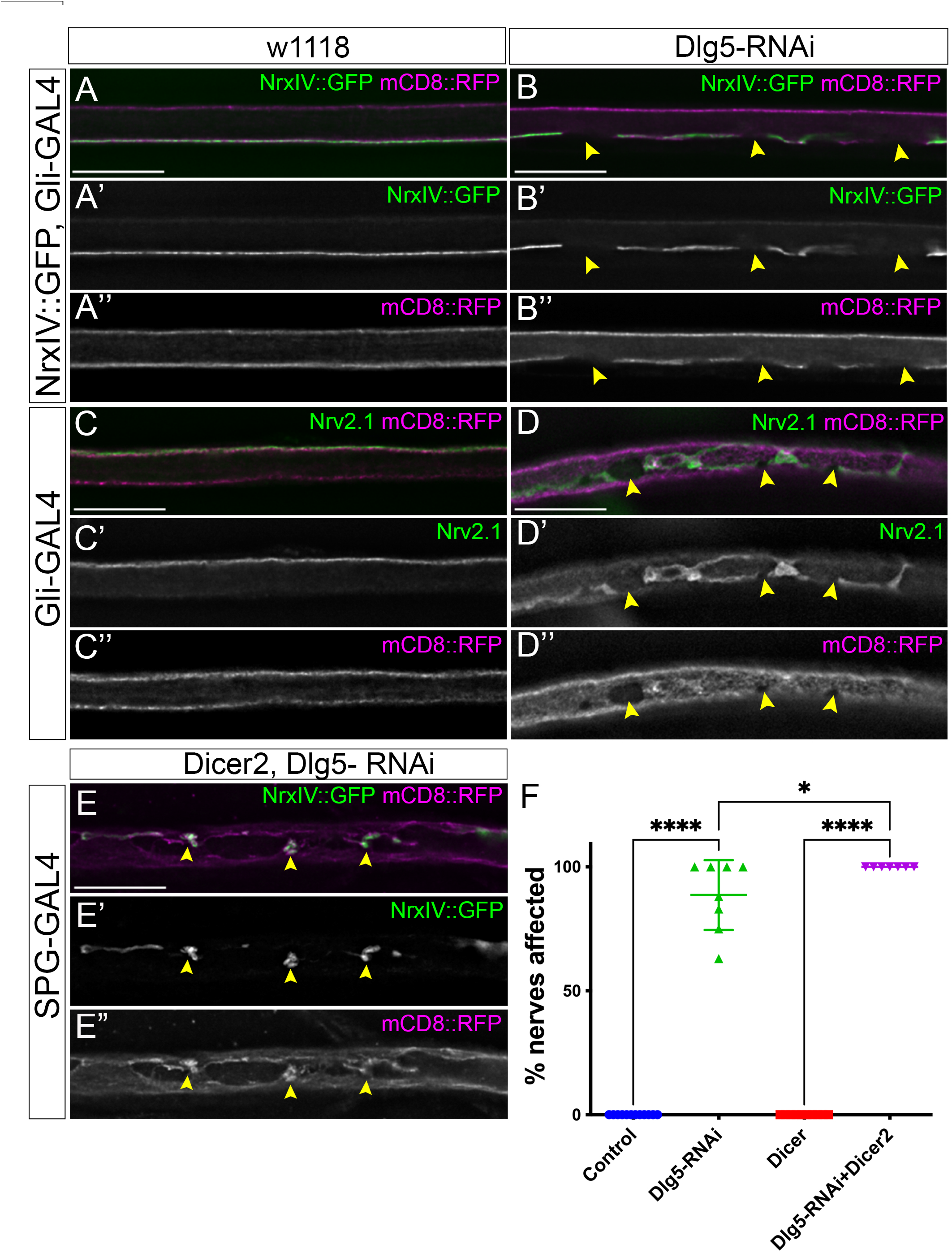
Knockdown of Dlg5 in the SPG disrupts septate junction and membrane morphology. The SPG drivers Gli-GAL4 (A-C) or SPG-GAL4 (E,F) driving mCD8::RFP (magenta) crossed to control (w[1118]) or Dlg5-RNAi. Scale bars: 15μm. **A-B:** Gli-GAL4 with NrxIV::GFP. Control (**A**) and Dlg5-RNAi (**B**) peripheral nerves with the SPG membrane labeled with mCD8::RFP (magenta) and SJs labelled with NrxIV::GFP (green). The SJs (**A’**) and SPG membranes (A”) were continuous. In *Dlg5-RNAi* nerves SJ strands were disrupted (B’), which corresponded to the disruptions in the SPG membrane (B” yellow arrowheads). **C-D:** Gli-GAL4. Control (**A**) and Dlg5-RNAi (**B**) peripheral nerves with the SPG membrane labeled with mCD8::RFP (magenta) and SJs immunolabelled for Nrv2.1 (green. The SJs (C’) and SPG membranes (C”) were continuous. In Dlg5-RNAi nerves SJ strands were disrupted (D’’), which corresponded to the disruptions in the SPG membrane (B” yellow arrowheads). **E:** SPG-GAL4, NrxIV::GFP. Dicer 2 enhanced the *Dlg5-RNAi phenotypes where* NrxIV and SJs formed accumulations (**E’**), which corresponded to accumulations and disruption in the SPG membrane (E” yellow arrowheads). **F:** Quantification of the SJ disruptions. The number of nerves per animal with disrupted SJs were quantified for SPG-GAL4 crossed to w[1118] (control) and Dlg5-RNAi or Dicer2 and Dicer2, Dlg5-RNAi. The mean and standard deviation are indicated. Statistical significance was determined by one-way ANOVA with Tukey’s *post hoc* multiple comparison test for each group (\**p* ≤0.5, **** *p* ≤0.0001)

### Dlg5 associates with Drosophila N-Cadherin and E-Cadherin

Prior studies have suggested that Dlg5 can interact in a number of cell types with β-catenin and cadherin family members. Specifically Dlg5 is found co-localized with cadherins (Liu et al., 2017), is involved in trafficking of cadherin-catenin adhesion complexes (Nechiporuk et al., 2007; Wang et al., 2014; Reilly et al., 2015), and links the vinexin-vinculin complex and beta-catenin at adherens junctions (Wakabayashi et al., 2003). We tested the possibility that Dlg5 also interacts with cadherins in peripheral glia. The presence and distribution of cadherins in Drosophila peripheral glia had not been established, so we began by looking for the presence of the two classical Drosophila cadherins; E-Cadherin (ECad; *shotgun*) and N-Cadherin (NCad; *CadN*) using genes endogenously tagged with mCherry or GFP (ECad::mCherry, ECad::GFP and NCad::mCherry (Venken et al., 2011)). Both NCad and ECad puncta were observed throughout the peripheral nerve (Fig. 4A,B) with both cadherins forming occasional short belt-like patterns in some parts of the nerve. More frequently, NCad and ECad were observed in puncta and we observed cadherin puncta throughout all glial layers (Fig. 4A-B, yellow arrows). The majority of the NCad (Fig. 4C’) and ECad (Fig. 4C”) puncta were distinct with minimal co-localization (Fig. 4C). The NCad and ECad puncta likely represent glial spot Adherens Junctions (sAJ) (Matzat et al., 2015) and these were observed between the three different glial layers with the majority occurring between the PG-SPG and SPG-WG processes (Fig. 4D, E)

**Figure 4.**
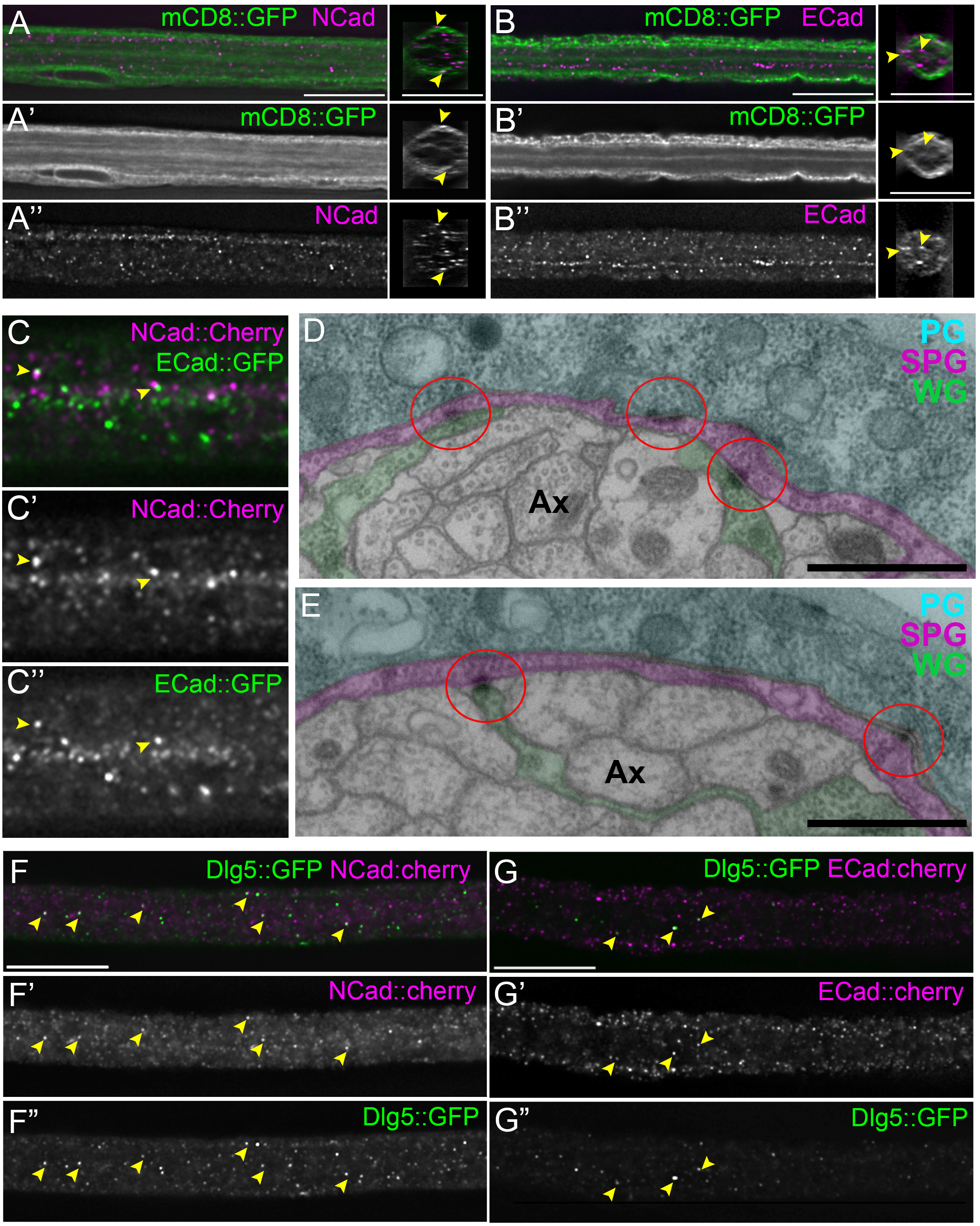
N-Cadherin and E-Cadherin are expressed in glial cells of the larval peripheral nerve. **A-B**: Control 3^rd^ instar peripheral nerves with NCad::mCherry (magenta, A,A”) and ECad::cherry (magenta, B,B”). The glial membranes were labeled with mCD8::GFP (green). Both Cadherins were distributed throughout the nerve indicated in the cross-sections of each peripheral nerve (yellow arrowheads). **C:** Longitudinal sections of peripheral nerves with NCad::mCherry (magenta, C,C’) and ECad::GFP (green, C,C”). Some but not all NCad overlaps with ECad puncta (yellow arrowheads). **D,E:** TEM analysis of control (repo-GAL4) nerve with the three glial layers falsed coloured: WG (green), SPG (magenta), PG (cyan). Spot adherens junctions are indicated (red circles). Scale bars: 1 μm **F,G:** Longitudinal sections of peripheral nerves with Dlg5::GFP (green) and NCad::mCherry (magenta, F,F’) and ECad::mCherry (magenta, G,G’). Some but not all Dlg5 overlaps NCad and ECad puncta (yellow arrowheads). Scale bars: 15μm

Next, we tested for Dlg5 and cadherin co-localization using Dlg5::GFP crossed with NCad::cherry (Fig. 4F) or ECad::cherry (Fig. 4G). Some but not all NCad (Fig. 4F’-F”) and ECad (Fig. 4G’-G”) puncta overlapped with Dlg5 in the peripheral nerve (yellow arrows). This partial overlap between NCad and Dlg5 is consistent with prior work in mouse embryonic fibroblasts, where Dlg5 only overlapped with a subset of NCad (Wang et al., 2014). In fibroblasts, Dlg5 is also partially associated with Rab11, a marker for recycling endosomes, as they fuse with cadherin-containing vesicles. To test whether Dlg5 in Drosophila peripheral glia are involved in trafficking pathways, we immunolabeled for a series of vesicle markers including Rab11, Rabsn5 (an early endosomal marker), Rab7 (a late endosomal maker), HRS (a multivesicular body (MVB) marker), Vps26 (a component of the retromer complex), Sec23 (a COPII vesicle marker), G130 (a Golgi maker) as well as syntaxin1A (a t-SNARE protein) (Supp Fig. 1). None of these subcellular markers consistently overlapped with Dlg5::GFP puncta in the peripheral nerve (Supp Fig. 1).

Dlg5 has been found associated with SJ proteins such as Dlg1 and Neurexin-IV (Nrx-IV) in Drosophila border cells (Liu et al., 2014a), yet in the peripheral glia Dlg5 did not localize with these proteins at the SJ. Therefore, we wanted to test to see if Dlg5 was associated with any other junctional complexes including gap junctions and focal adhesions (Supp Fig. 2). However, we did not observe any co-localization between the Dlg5::GFP and gap junctions immunolabeled with Inx2 (Supp Fig. 2C-C”) nor with focal adhesions, immunolabeled for the integrin beta-subunit βPS (Supp Fig 2D-D”). Therefore, Dlg5 is unlikely to be part of the gap junction or focal adhesion complex. In addition, pan-glial knockdown of Dlg5 had no effect on Inx2 labeled gap junctions, or βPS labeled focal adhesions. Overall, our results suggest that of the potential junctional domains in peripheral glia, Dlg5 is the most frequently associated with cadherins. However, this association appears to be limited to a subset of NCad and ECad.

### E-Cadherin is required in glial cells of larval peripheral nerve

To determine whether Dlg5 and cadherins work in a similar pathway, we next examined the function of NCad and ECad in the peripheral nerve. To assay for abnormalities in glial morphology, *repo-GAL4* was used to express RNAi and label the glial membranes with mCD8::GFP and immunolabeling for Dlg1 used to assess SJ morphology. Normal glial morphology and Dlg1 distribution was observed in all the control nerves (*repo>mCD8::GFP, Dicer2*) (n=6 larvae) (Fig. 5A-A”). Pan-glial knockdown of ECad (*repo>ECad-RNAi, Dicer2*) using multiple RNAi lines resulted in disrupted glial membranes and the appearance of “hole-like” structures within the glial membrane in 71.4% nerves of the strongest RNAi (27082; n= 4 larvae) (Fig. 5B’). Dlg1 was mislocalized in 72% of the affected nerves (n=4 larvae) (Fig. 5B”). Surprisingly, knockdown of NCad (*repo>NCad-RNAi, Dicer2*) using multiple RNAi lines did not affect glial morphology or Dlg1 distribution (data not shown).

**Figure 5.**
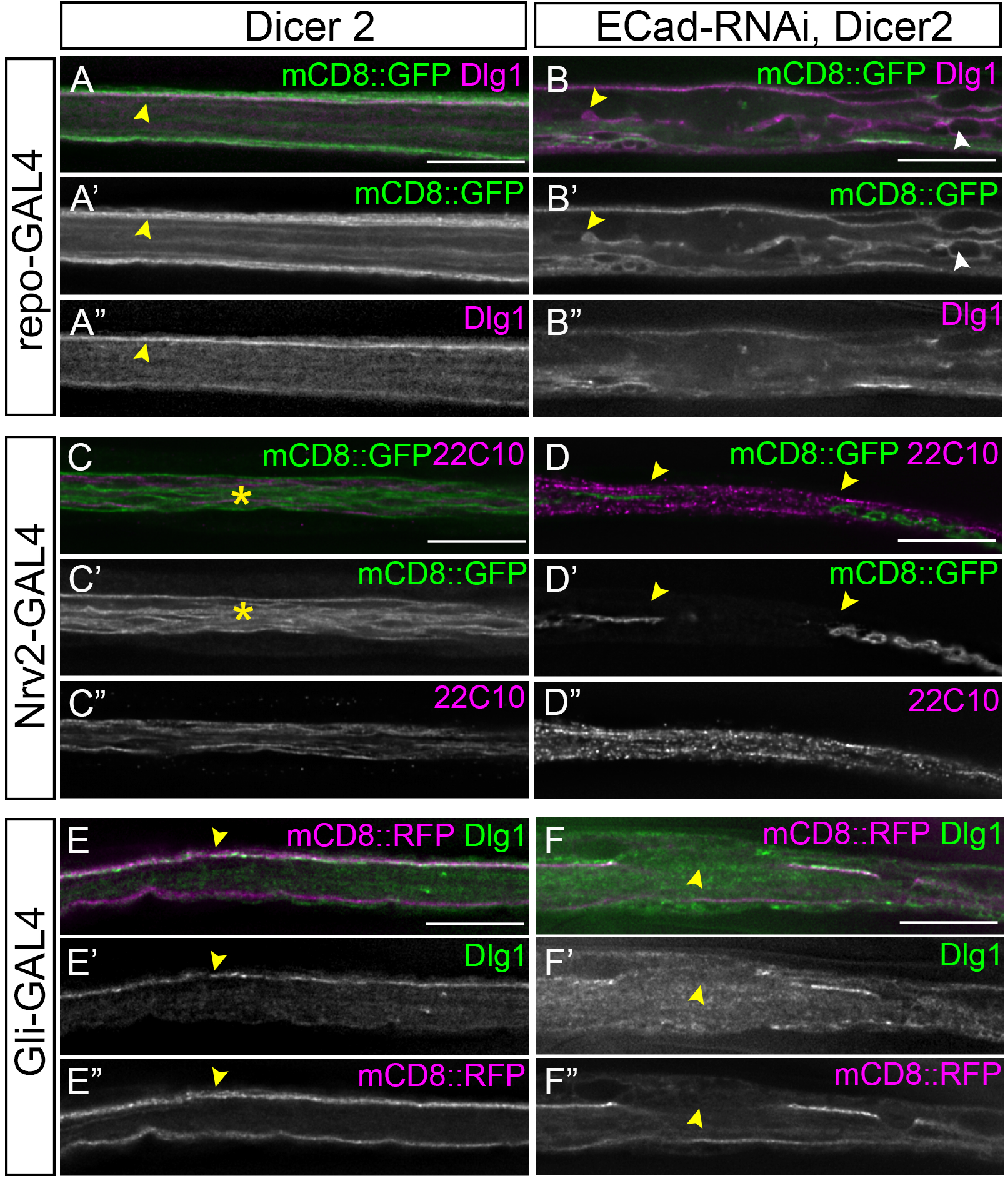
Knockdown of ECad in the WG and SPG affects glial morphology. **A-B**: All glia. repo-GAL4 crossed to Dicer2 (control) (A) and ECad-RNAi, Dicer2 (B) Peripheral nerves with all glial membranes labeled with mCD8::GFP (green). SJ were immunolabeled with Dlg1 (magenta). Glial membranes (A’) and a linear SJ (A”) extends along the length of the nerve in control. With ECad-RNAi, glial membranes are disrupted (yellow arrowheads, B,B’) and vacuole-like or hole-like structures observed (white arrowheads, B’B”) and the SJ disorganized. **C-D**: Wrapping glia. Nrv2-GAL4 crossed to Dicer2 (control)(C) and ECad, Dicer2 (D). Longitudinal sections of peripheral nerves with WG membranes labeled with mCD8::GFP (green) and axons immunolabeled with anti-Futsch (22C10, magenta). The WG membranes continuously extend along the length of control (green, C,C’). With ECad-RNAi, WG membranes were discontinuous and disrupted (yellow arrowheads, D,D’). The axon morphology in ECad-RNAi peripheral nerves looked similar to controls (magenta, D,D’’). **E-F**: Subperineurial glia. Gli-GAL4 crossed to Dicer 2 (control)(E) and ECad-RNAi, Dicer2 (F) with the SPG membrane labeled with mCD8::RFP (magenta) and the SJ immunolabeled with Dlg1 (green). The SPG membrane is continuous along the length of control (magenta, E”) and the SJ forms a continuous line (E’). In ECad-RNAi nerves the membrane was discontinuous and disrupted (yellow arrowheads, F”) and whereas Dlg1 was lost in regions of the nerve where the SPG membrane was disrupted. Scale bars: 15μm

To determine if ECad is required in one or all glial layers, we knocked down ECad in individual glial layers using the strong ECad-RNAi line (27082). ECad knockdown in the wrapping glia using Nrv2-GAL4 paired with Dicer2 resulted in fewer and discontinuous wrapping glia strands in 89% of nerves (*Nrv2>ECad-RNAi, Dicer2*) (n=4 larvae) (Fig. 5D). Axon morphology was not affected by the knockdown of ECad (Fig. 5D”). Control nerves (*Nrv2>Dicer2*) had the expected multiple, continuous wrapping glia strands ensheathing the peripheral axons (Fig. 5C). Therefore, ECad is required in the wrapping glia and is necessary for axon ensheathment. Similar to the pan-glial knockdown of NCad, knockdown in the wrapping glia did not affect glial morphology and the morphology of the axons was comparable to the control. We next tested the function of ECad in the SPG layer using the SPG driver, Gli-GAL4, to drive RNAi and label the SPG membranes with mCD8::RFP. The SPG membranes observed in all control nerves (*Gli>mCD8::RFP, Dicer2*) were continuous and Dlg1 was correctly localized to the septate junction (n= 6 larvae) (Fig. 5E). In comparison knockdown of ECad in the SPG (*Gli>ECad-RNAi, Dicer2*) resulted in nerves with discontinuous SPG membranes (Fig. 5F) (n=6 larvae). The gaps in the SPG membrane coincided with the loss of Dlg1 (Fig. 5F’,F”; yellow arrows). Therefore, ECad is required in the SPG for proper ensheathment and SJ formation while the knockdown of NCad had no effect.

ECad knockdown affected SPG morphology and resulted in SPG defects similar to those observed with the knockdown of Dlg5. To further characterize and quantify these phenotypes, we used NrxIV::GFP as a marker to assess any changes in the SJ when E-Cad was knocked down in the SPG (*moody>mCD8::RFP, ECad-RNAi*). Similar to the loss of Dlg5, ECad knockdown results in gaps or breaks and aggregates of NrxIV::GFP within the SJ strands in 48% of nerves (Fig. 6C-C’) along with disruption to the glial membrane (Fig. 6C”,D”)(n=6 larvae). Increasing the efficiency of the knockdown with Dicer2 (*moody>mCD8::RFP, ECad-RNAi, Dicer2*) strengthened the phenotypes to the point where NrxIV::GFP was frequently observed in aggregates (Fig. 6E-E’,F-F’; yellow arrows) and the SPG membrane appeared collapsed (Fig. 6E”,F”). The SJ strands in controls (*moody>mCD8::RFP, Dicer2*) were continuous and NrxIV::GFP aggregates were never observed (Fig. 6A-A’) (n=10 larvae). The presence of gaps or disruption to the SJ was quantified and the percentage of nerves with SJ gaps was significantly higher compared to controls (Fig. 6B).

**Figure 6.**
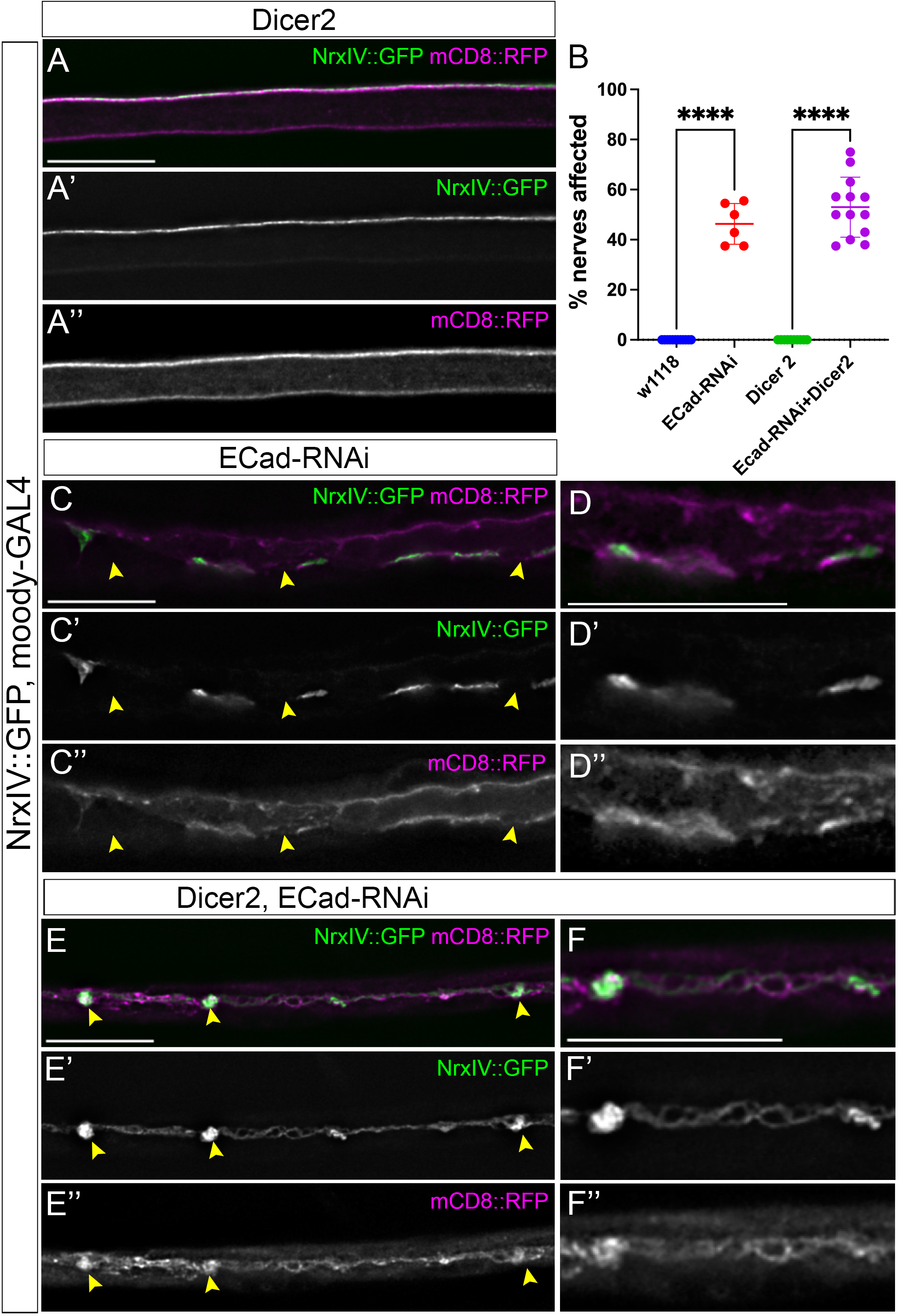
Knockdown of ECad in the SPG disrupts the septate junctions. Longitudinal sections of 3^rd^ instar peripheral nerves with the SPG driver, moody-GAL4 driving mCD8::RFP (magenta) to label the glial membranes and with NrxIV::GFP (green) to mark the SJ. Scale bars: 15μm **A:** Control. Moody-GAL4 crossed to w[1118]. Glial membranes (A”) and a linear SJ (A’) extends along the length of the nerve in control. **B:** Quantification of the ECad-RNAi effects on the SJ. The percentage of nerves affected per larvae were determined for each genotype. The mean and standard deviation are indictated. Statistical significance was determined by one-way ANOVA with Tukey’s *post hoc* multiple comparison test for each RNAi compared to the control (**** *p* ≤0.0001). **C-D**: ECad-RNAi. Moody-GAL4 crossed with ECad-RNAi. The SJ and glial membranes contained breaks or holes in the immunolabeling (yellow arrowheads). D represents a digital magnification (200%) of C to highlight the membrane and SJ breaks. **E-F**: ECad-RNAi plus Dicer2. Moody-GAL4 crossed with ECad-RNAi and Dicer2 to enhance the RNAi efficiency. NrxIV formed aggregates in the membrane which appeared collapsed (yellow arrowheads). F represents a digital magnification (200%) of E to highlight the NrxIV aggregates.

### N-Cadherins play redundant roles in the larval peripheral nerve

Although NCad is expressed in the peripheral nerve, knockdown of NCad did not affect glial morphology. There are two NCad loci in *Drosophila*, CadN (N-Cad) and a duplication CadN2 (N-Cad2) (Prakash et al., 2005). CadN2 shares 72.5% amino acid sequence identity and is partially redundant to NCad (Yonekura et al., 2007). Therefore, it is possible that the loss of N-Cad is compensated by CadN2. We knocked down CadN2 in all glia using multiple RNAi lines (*repo>CadN2-RNAi*) and similar to N-Cad knockdown, CadN2 knockdown did not affect glial morphology. To test for redundancy between these genes, we knocked down both NCad and CadN2 in the SPG (*moody>NCad-RNAi, CadN2-RNAi*) and assayed for *NrxIV::GFP* labeled SJs. We found that reducing the levels of both NCad and CadN2 resulted in defects in 35% of the nerves (Fig. 7B-B’’) (n=8 larvae). When we quantified the number of nerves that contained gaps or breaks in the SJ, 13.5% of nerves were affected (Fig. 7E), which was significantly greater than either RNAi alone or control.

**Figure 7.**
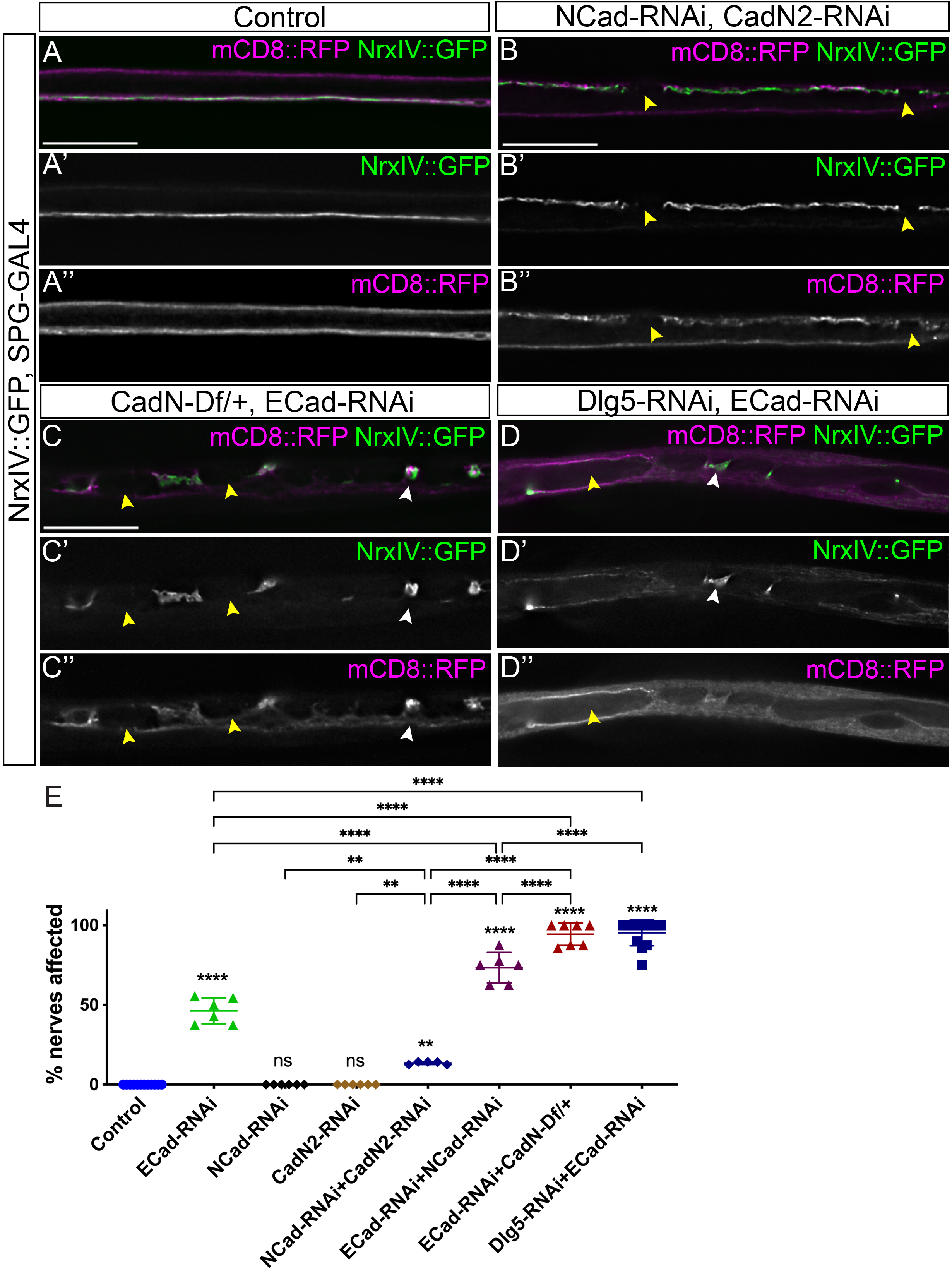
Knockdown of Dlg5, ECad and NCad disrupts septate junctions. Longitudinal sections of 3^rd^ instar peripheral nerves with the SPG driver, moody-GAL4 driving mCD8::RFP (magenta) to label the glial membranes and with NrxIV::GFP (green) to mark the SJ. Scale bars: 15μm **A:** Control. Moody-GAL4 crossed to w[1118]. Glial membranes (A”) and a linear SJ (A’) extends along the length of the nerve in control. **B**: NCad-RNAi, CadN2-RNAi. Moody-GAL4 crossed with NCad-RNAi, CadN2-RNAi. The SJ (B’) and glial membranes (B”) contained breaks or holes (yellow arrowheads). **C**: ECad-RNAi, CadN-Df/+. Moody-GAL4 crossed with ECad-RNAi and a deficiency that uncovers both NCad and CadN2. NrxIV formed aggregates (C’) and those corresponded to membrane concentrations (white arrowhead). Disrupted of the SJ occurred in the same places as loss or holes in the glial membrane (yellow arrowheads). **D:** Dlg5-RNAi, ECad-RNAi. Moody-GAL4 crossed with Dlg5 and ECad-RNAi. NrxIV formed aggregates (C’) and those corresponded to membrane concentrations (white arrowhead). SJ were reduced and large holes were observed in the glial membranes (yellow arrowheads). **E:** Quantification of the RNAi effects on the SJ. The percentage of nerves affected per larvae were determined for each genotype. The mean and standard deviation are indicated. Statistical significance was determined by one-way ANOVA with Tukey’s *post hoc* multiple comparison test for each RNAi compared across multiple genotypes. p values compared to control are indicated above each genotype. (\**p* ≤0.5, \*\**p* ≤0.01, *** *p* ≤0.001, **** *p* ≤0.0001, non-significant differences are not indicated)

These defects were similar to those we observed with the knockdown of ECad in the SPG suggesting that NCad and CadN2 are not only redundant with each other but also perform some of the same functions as ECad. To test if ECad and NCad are redundant, we performed a double knockdown of NCad and ECad in the SPG (*moody>NCad-RNAi, ECad-RNAi)*. Knockdown of NCad significantly enhanced the ECad-RNAi phenotypes compared to either RNAi line alone (n=6 larvae) (Fig. 7E). To further test if all three cadherins perform similar functions in the peripheral glia, we knocked down ECad in conjunction with a heterozygous deficiency (*Df(2L)BSC148)* that deletes both NCad and CadN2 (*CadN[Df]/+; moody>ECad-RNAi*). The reduction of both NCad genes strongly enhanced the ECad RNAi phenotypes (Fig. 7C-C”) (n=10 larvae) with gaps or breaks within the SJ domain (yellow arrowheads) and aggregates of the NrxIV::GFP protein (white arrowheads). The penetrance of the gaps or breaks present within the SJs was significantly greater compared to control, ECad-RNAi alone or the ECad+NCad double RNAi combination (Fig 7E). In summary these results suggest that ECad, NCad and CadN2 are partially redundant in the peripheral glia.

*Dlg5 and cadherins function in the same pathway and underlie SJ and spot AJ formation* The knockdown of ECad alone or in combination with loss of NCad specifically in the SPG phenocopied the SJ phenotypes observed with the knockdown of Dlg5 and this led to the hypothesis that these proteins might function in the same or parallel pathways. To test this, we combined Dlg5 and ECad RNAi lines and assessed the effects of simultaneous knockdown on SJ formation. We observed a strong enhancement of the ECad-RNAi phenotypes by Dlg5-RNAi (Fig. 7D-D”) (n=13 larvae) where the percentage of nerves affected with Dlg5+ECad-RNAi were significantly greater than ECad-RNAi alone (n=6 larvae) and ECad+NCad-RNAi (Fig. 7E). The effects of Dlg5+ECad-RNAi were on par with the severity and penetrance of we observed with ECad-RNAi plus the heterozygous NCad+CadN2 deficiency. These results suggest that Dlg5 and cadherin function synergistically in the SPG and that Dlg5 may play a role in SJ formation.

We attempted to test the effects of Dlg5 knockdown on the formation of sAJ using immunolabeling with ECad::cherry and NCad::cherry, however this analysis was difficult due to the distribution of cadherin puncta within all three glial layers. To better assess the effects on sAJ, we used TEM analysis of Dlg5-RNAi knockdown using the pan-glial driver (*repo>Dlg5-RNAi, Dicer2*) and assessed for the presence of sAJ (Fig. 8). In controls (*repo>Dicer2*)(n=3 larvae) we observed sAJ between the multiple glial layers, specifically between the PG and SPG (yellow) and between SPG and WG (green) with an average of 1.8 sAJ per section for each (Fig. 8E; red circles). We observed rare instances of sAJ between WG processes and found sAJ between SPG only when two SPG where present (Fig. 8E). Knockdown of Dlg5 (*repo>Dlg5-RNAi*) lead to a significant reduction in the sAJ between both the PG-SPG and the SPG-WG membranes (on average 0.3 and 0.5 per section respectively) (Fig. 8A,B) (n=2 larvae). This was similar to the loss of sAJ between the glial layers when ECad-RNAi was expressed (*repo>ECad-RNAi, Dicer2*) (Fig. 8C,D) where the presence of sAJ decreased to an average of 0.1 between PG-SPG and 0.3 between SPG-WG per section (n=5 larvae). Our data suggest that knockdown of Dlg5 or ECad have the same effects resulting in a reduction in sAJs junctions between the glial layers.

**Figure 8.**
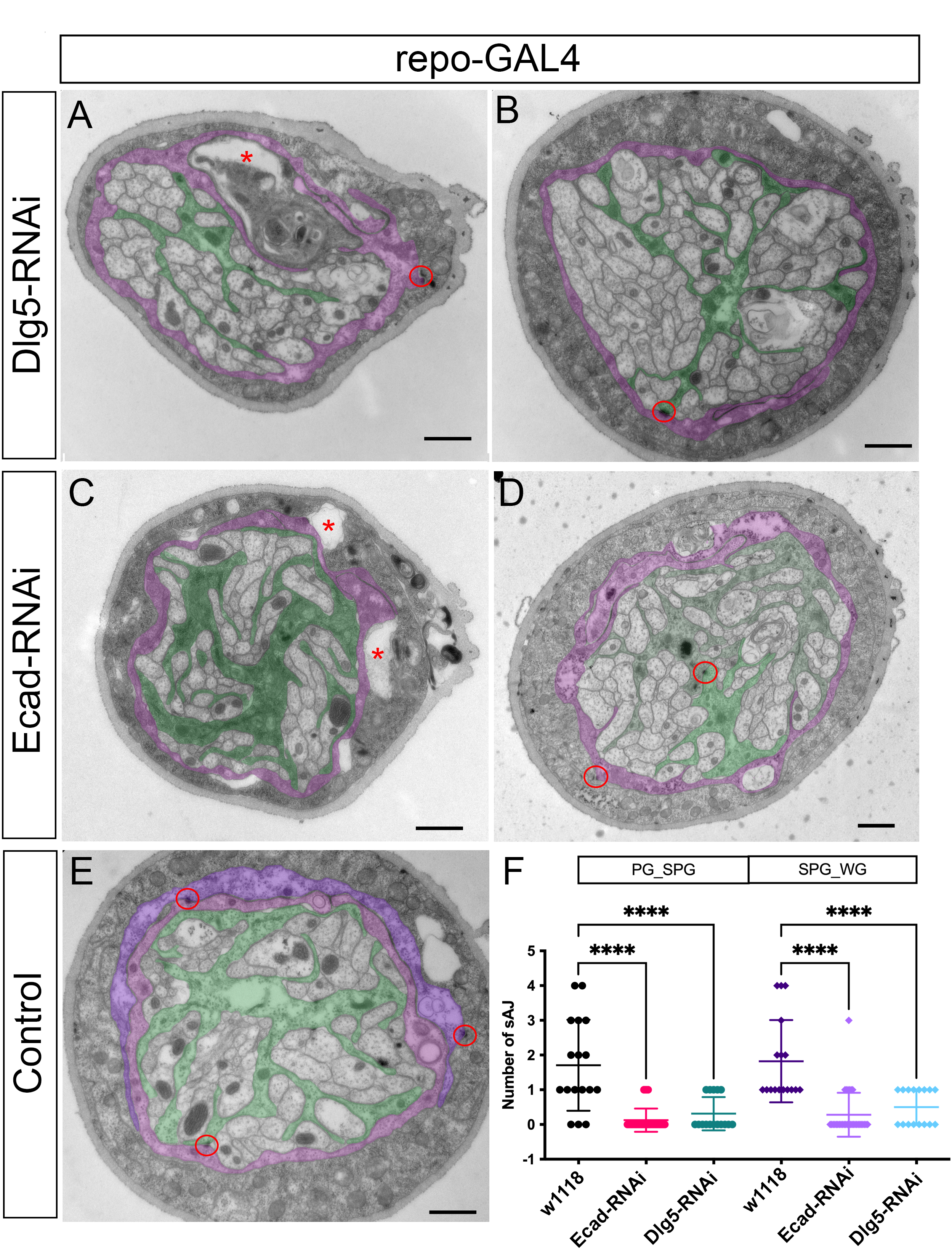
Knockdown of Dlg5 and ECad leads to loss of spot adherens junctions. Transmission electron microscope images of repo-GAL crossed to Dlg5-RNAi (A-B), ECad-RNAi (C-D) or w1118 (E) with the SPG and WG layers false coloured (SPG: magenta; WG: green). The outer PG layer and the inner axons are not coloured. Spot adherens junctions are indicated with a red circle. Unusual gaps within the SPG or between the SPG and WG are indicated with an asterisk (red). Scale bars: 1 μm. F: Quantification of the number of spot adherens junctions identified between the PG and SPG membranes and the SPG and WG membranes. The mean and standard deviation are indicated. Statistical significance was determined by one-way ANOVA with Tukey’s *post hoc* multiple comparison test for each group compared to control (**** *p* ≤0.0001)

#### Loss of Dlg5 and ECad disrupts the blood-brain barrier and larval locomotion

Glial septate junctions form the blood-barrier in the Drosophila CNS and PNS. Given the strong effects of loss of both Dlg5 and the Cadherins on the septate junctions, we next tested for changes to the blood-brain barrier using a dextran dye assay (Fig. 9). Dye was blocked, as expected, from entry into the brain lobes and ventral nerve cord of the CNS in controls (*moody-GAL4*) (Fig. 9A) (n=30 larvae), while knockdown of NrxIV as a positive control (*moody>NrxIV-RNAi*) disrupted the SJ and led to significant dye penetration (Fig. 9B, F) (n=2 larvae). Knockdown of ECad but not Dlg5 lead to a small but significant increase in the penetration of dye into the CNS (Fig. 9C,D,F) (n=72 and 53 larvae respectively) and the double knockdown of Dlg5+ECad (*moody>Dlg5-RNAi, ECad-RNAi*) (n=70 larvae) had significantly more dye penetration compared to either RNAi alone (Fig. 9E,F).

**Figure 9:**
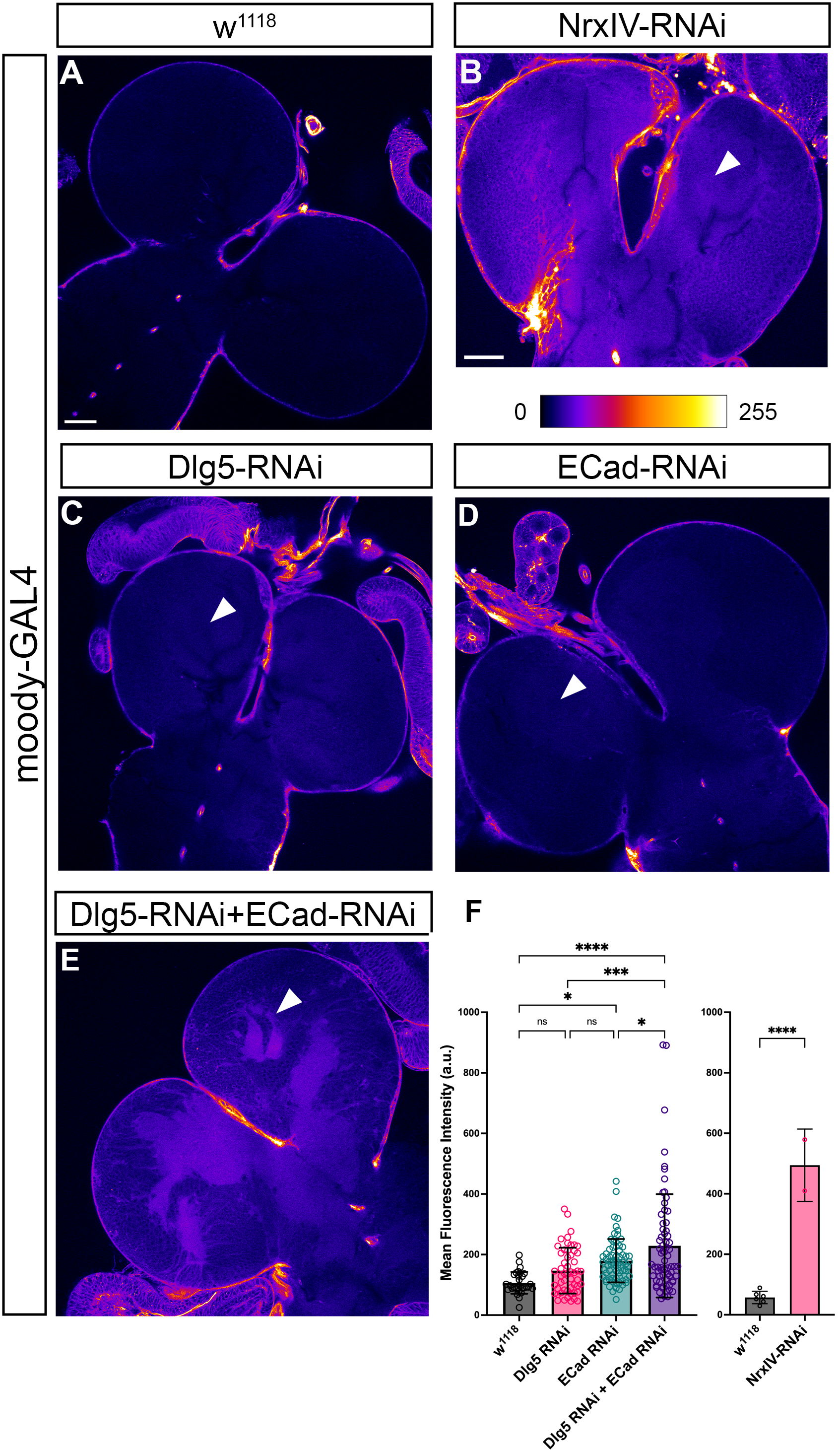
Knockdown of Dlg5 and ECad impairs blood-brain barrier integrity. **A-D :** Fire LUT showing blood-brain permeability assay using moody-GAL4>mCD8::GFP crossed to w[1118] (control)(A), Dlg5-RNAi (B), ECad-RNAi (C), Dlg5-RNAi, ECad-RNAi (D) or NrxIV-RNAi (E). White arrows highlight regions of significant dye labeling of the neuropile. Scale bars: 150 μm. **F:** Quantification of mean fluorescent intensity inside the brain lobe. The mean and standard deviation are indicated. Dye penetration for NrxIV-RNAi was determined at a lower laser intensity than the other genotypes. Statistical significance was determined by one-way ANOVA with Tukey’s *post hoc* multiple comparison test for each group (\**p* ≤0.5, \*\**p* ≤0.01, *** *p* ≤0.001, **** *p* ≤0.0001, non-significant differences are not indicated).

We wanted to determine if the changes to the glial layers, the septate junctions and the disruption of the blood-brain barrier affected nervous system function. To this end, we measured various parameters of locomotor behavior (Fig. 10) including the total distance and the average speed at which 3^rd^ instar larvae travelled. We found that in controls (*moody-GAL4* crossed to *w[1118]*), larvae travelled on average a total distance of 67mm at an average speed of 1.12 mm/sec (Fig. 10A, F-G). We used the knockdown of NrxIV (*moody>NrxIV-RNAi*), which disrupts SJs and the blood-barrier as a positive control and larvae motion was significantly compromised (Fig. 10E-G). When compared to control larvae, both parameters were significantly reduced after ECad knockdown (*moody>ECad-RNAi*)(Fig. 10C,F,G) and Dlg5 knockdown (*moody>Dlg5-RNAi*)(Fig.10B,F,G). The combined knockdown of Dlg5 and ECad (*moody>ECad-RNAi, Dlg5-RNAi*) significantly reduced locomotion (Fig. 10D,F,G) compared to ECad-RNAi or Dlg5-RNAi alone and matched the disruption in locomotion observed with NrxIV-RNAi. Our results suggest that the disruption of the SJ domains leads to changes in the integrity of the blood-brain barrier and to changes in animal locomotion.

**Figure 10.**
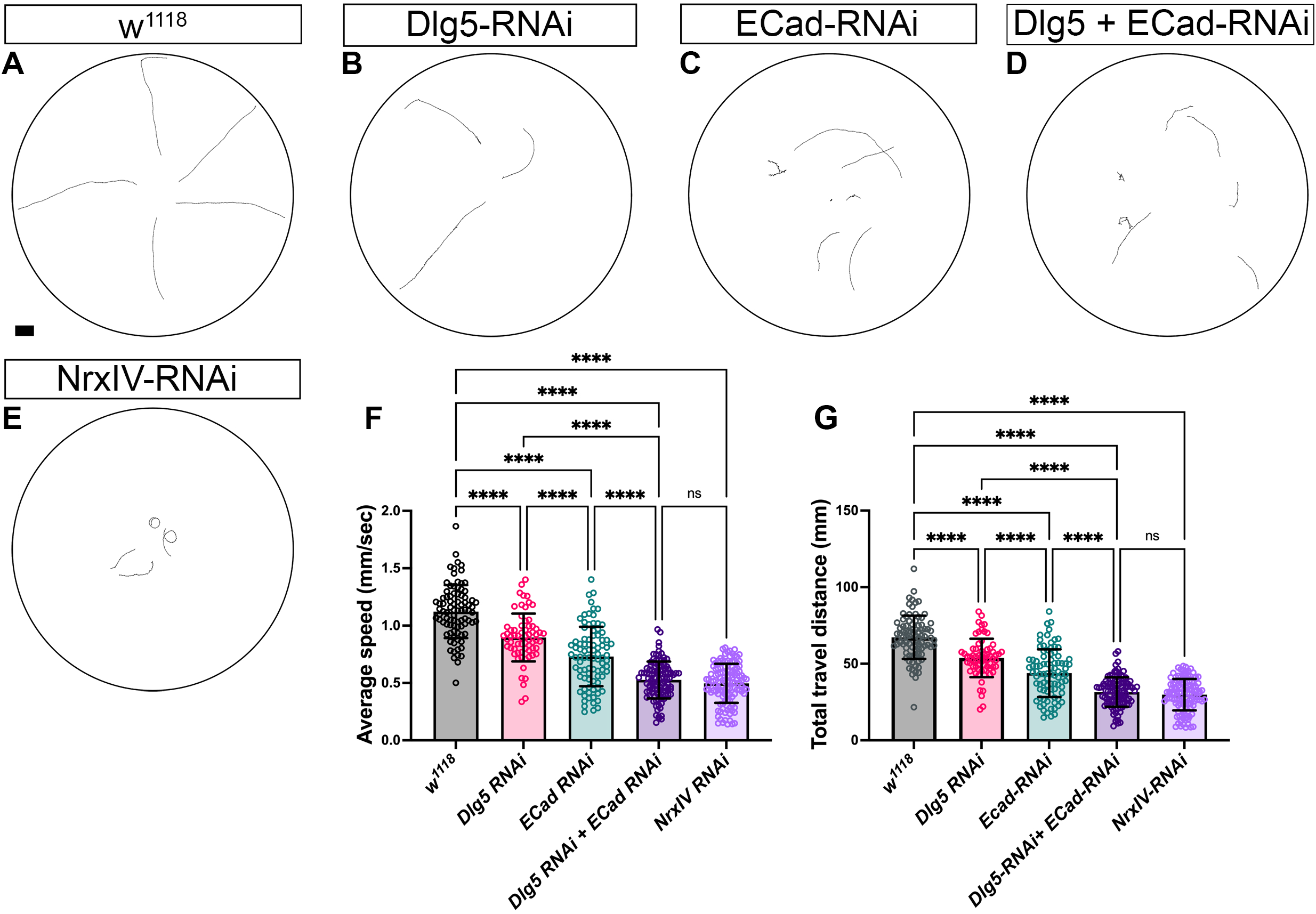
Loss of Dlg5 and ECad in the SPG impairs animal locomotion. **A-E** : Movement trajectories of 3rd instar larvae from moody-GAL4 flies crossed with w[1118] (A), Dlg5-RNAi (B), ECad-RNAi (C), Dlg5-RNAi, ECad-RNAi (D), and NrxIV-RNAi (E). Scale bars: 10 mm. **F-G**: Comparison of average speed (F) and total travel distance (G) of moody-GAL4 driving Dlg5-RNAi, ECad-RNAi, Dlg5+ECad-RNAi, NrxIV-RNAi and control, w[1118]. The mean and standard deviation are indicated. Statistical significance was determined by one-way ANOVA with Tukey’s *post hoc* multiple comparison test for each group (\**p* ≤0.5, \*\**p* ≤0.01, *** *p* ≤0.001, **** *p* ≤0.0001, non-significant differences are not indicated).

## Discussion

Our aim was to discover PDZ domain scaffolding proteins that play a role in glial ensheathment of peripheral axons. We identified that the MAGUK protein Discs-large 5 (Dlg5) has a role in subperineurial glia and septate junction morphology. To investigate the role of Dlg5 further we focused on cadherins and found that loss of N-Cadherin and E-Cadherin phenocopied the loss of Dlg5 leading to gaps in the SPG and septate junctions. This leads to a model where Dlg5 plays a role in conjunction with Cadherins in glial membrane stabilization and SJ formation in the subperineurial glia.

Dlg5 performs multiple independent functions in both mammals and Drosophila including regulation of cell proliferation, polarity, growth, and trafficking of junctional complexes and these functions are highly context specific (Kwan et al., 2016; Liu et al., 2017; Venugopal et al., 2020). One established function for Dlg5 is a role in polarity. In Drosophila, Dlg5 plays a role in establishing apical-basal polarity in follicular epithelia, where Dlg5 localizes at the apical membrane and adherens junction in early egg chambers and where loss of Dlg5 generates epithelia polarity defects (Luo et al., 2016). In follicular epithelia, Dlg5 promotes the apical membrane localization of Crumbs to establish polarity. Our data suggests that in the context of glial cells, Dlg5 also may play a role in maintaining cell polarity as we observed changes to the SJ and the distribution of Dlg and Scrib. However, unlike the follicular epithelial cells, Dlg5 most likely does not function through changes to Crumbs in the SPG as peripheral glia do not express Crumbs and knockdown of Crumbs had no effect. Glial cells are likely polarized by a different mechanism in a Crumbs independent manner. We also failed to detect Par-3 or Par-6 in the peripheral glia and RNAi knockdown of these apical polarity proteins had no effect. The unconventional polarization mechanism adopted by glia could be attributed to differences in the architecture of glial cells compared to epithelia. We did find that Sdt, a component of the Crumbs complex, is required in the peripheral glia, which raises the question as to whether Sdt interacts with non-canonical partners to perform similar functions as seen in epithelial cells or whether Sdt might have an entirely different and novel function in glial cells. The latter is more likely as Sdt knockdown does not affect SJ morphology.

Another potential Dlg5 function could be through the interface between Cadherin function and the glial cytoskeleton or in the assembly of spot adherens junctions. Dlg5 links the vinexin-vinculin complex and beta-catenin at sites of cell-cell contact in mammalian fibroblasts and epithelial cells (Wakabayashi et al., 2003). Loss of Dlg5 in the SPG resulted in the reduction in the number of spot adherens junctions and enhanced the SPG defects generated by loss of Cadherins. However, the degree of overlap between Dlg5 and Cadherins in the 3^rd^ instar larvae was not extensive suggesting a role of Dlg5 in the construction of the sAJs rather than the maintenance and ongoing stability of these junctions. This mirrors prior observations where Dlg5 was necessary for the trafficking of N-cad/beta-catenin complexes to the membrane (Nechiporuk et al., 2007).

It is possible that Dlg5 function with regards to junctional complexes is highly context specific. In the Drosophila ovary, Dlg5 is found associated with basolateral SJ proteins (including Dlg1, Neuroglian, ATPα and Lachesin) and to a smaller extent the AJ in border cells (Luo et al., 2019), and Dlg5 was localized with Dlg1 in stage 10 follicular epithelia (Luo et al., 2016). The loss of Dlg5 disrupted the distribution of Arm (beta-catenin) in border cells (Luo et al., 2019) and E-Cadherin in follicular cells (Luo et al., 2016), yet had no effect on the distribution of Dlg1 or the SJ protein Neuroglian (Luo et al., 2019). In wing discs, Dlg5 mutants have a reduction of apical domain determinants but not a loss of cell polarity and a very specific defect in the recruitment of N-Cadherin to adherens junctions but not E-Cadherin (Venugopal et al., 2020). Our results shown that in the SPG, Dlg5 was not found associated with any junctional complexes except with puncta of NCad and ECad, and was excluded from the SJ. As well loss of Dlg5 resulted in the disruption of both sAJ and SJ morphology suggesting a global defect in cell polarity rather than a specific disruption to a single junctional domain.

A third potential function for Dlg5 is in controlling cell proliferation (Nakamura et al., 1998; Liu et al., 2017) where Dlg5-knockdown cells grow faster. Dlg5 function appears to be highly cell specific as Dlg5 can function either as tumour suppressor or tumour enhancer in different cancer cell lines (Andolfi et al., 2022) (Liu et al., 2014b). As SPG are polyploid cells and do not undergo mitosis, we do not think the role of Dlg5 is to control SPG proliferation. However, the SPG are very large cells and polyploidy is necessary for glial growth to match the expanding CNS, and required for a functional blood–brain barrier (Unhavaithaya and Orr-Weaver, 2012). A key component in the SPG size increase is the Hippo pathway (Liu et al., 2017), a key regulator of organ size and tissue homeostasis in many tissues (Limmer et al., 2014; Yu et al., 2015; Li and Guan, 2022). The Hippo pathway regulates the transcriptional regulator Yorkie specifically by inactivating Yorkie through phosphorylation and degradation (Comoletti et al., 2003). In Drosophila, Yorkie is necessary for normal glial cell numbers and expansion (Reddy and Irvine, 2011), and for the extensive growth of SPG (Li et al., 2017). Loss of Yorkie activity in the SPG causes abnormal endoreplication, defective septate junctions and BBB disruption (Li et al., 2017). There is evidence that Dlg5 can indirectly regulate Hippo signaling in multiple Drosophila tissues, where loss of Dlg5 leads to activation of Hippo signaling and the repression of Yorkie (Kwan et al., 2016; Kaya-Copur et al., 2021; Venugopal et al., 2020). Our results found that loss of Dlg5 and Cadherins leads to SJ and BBB disruption suggesting that, if Hippo is involved, in the SPG loss of Dlg5 leads to a decrease in Yorkie activity and an increase in Hippo signaling. Given that the polypoid SPG do not divide but rather grow in size, it is possible that loss of Dlg5 leads to reduced growth in SPG which manifests as hole-like structures in the SPG and the SJ. However, whether Dlg5 regulates SPG and SJ development through Hippo pathway or other pathways remains to be elucidated.

## ACKNOWLEDGMENTS

This work was supported by UBC Bioimaging facility (RRID: SCR_021304). Stocks were obtained from the Bloomington Drosophila Stock Center (NIH P40OD018537) and the Vienna Drosophila Resource Center (VDRC, www.vdrc.at). This research was funded by a grant from the Natural Sciences and Engineering Research Council of Canada (NSERC) and the Canadian Institutes of Health Research (CIHR).

