## Supplementary material for "Dlg5 and Cadherins are key to peripheral glia integrity": Supplmental Figures

**Supplemental Data:**

**Supplemental Table 1**: Summary of PDZ proteins screened
The stock numbers of each UAS-PDZ-RNAi line obtained from the Bloomington Drosophila Stock Center (BDSC) or the Vienna Drosophila Research Center (VDRC) are provided. The glial phenotypes observed by expressing each RNAi line in glial cells are described. Candidates were divided into: positive hits (yellow), those tested using only one RNAi line (orange) and those candidates for which two RNAi lines were tested and one resulted in a glial phenotype whereas the other did not (green). The RNAi lines that resulted in glial phenotypes are highlighted in purple.

**Table 1 legend**

positive hits, more than 1 RNAi lead to a glial phenotype

only 1 RNAi tested

RNAis that resulted in phenotypes

candidates for which 1 RNAi lead to a phenotype while the other did not

**Supplemental Figure 1:**
**A:** Longitudinal sections of peripheral nerves labeled with Dlg5 and Vps26.
The retromer complex was immunolabeled with Vps26 (magenta, A') and Dlg5 was labeled with Dlg5::GFP (green, A''). The Dlg5 puncta (green, A) only overlapped with a small subset of Vps26 (yellow arrowheads).
**B:** Longitudinal sections of peripheral nerves labeled with Dlg5 and Recycling endosomes. Rab 11-positive recycling endosomes was labeled with Rab-11 antibody (magenta, B') and Dlg5 was labeled with Dlg5::GFP (green, B''). The Dlg5 puncta (green, A) only overlapped with a small subset of Rab-11 (yellow arrowheads).
**C-G:** Longitudinal sections of peripheral nerves labeled with Dlg5, Golgi (C), late endosomes (D), multivesicular bodies(E), t-SNAREs (F) and COPII vesicles (G). The Golgi complex was immunolabeled with G130 (C), late endosomes were marked using an anti-Rab-7 antibody (D), multivesicular bodies was immuno-labelled with HRS (E), t-SNAREs were visualized using the anti-sytx1A antibody (F), and COPII vesicles were immuno-labelled with anti-Sec23 antibody (G). None of the markers shows any overlaps with Dlg5. Scale bars = 15µm.'

**Supplemental Figure 2:**
Longitudinal sections of peripheral nerves labeled with Dlg5 and Inx2 immunolabeled gap junction complexes in the peripheral nerve. (A'', green) in the peripheral nerve. Dlg5 puncta were present throughout the nerve but did not co-localize with Inx2 labeled gap junction plaques (A', magenta).
**A:** Cross-sections of the peripheral nerve showing that Dlg5 does not co-localize with Inx2.
**B:** Longitudinal sections of peripheral nerves labeled with Dlg5 and focal adhesion complexes immunolabeled with the beta subunit of integrin, myospheriod (βPS). Dlg5 puncta (green, B") were observed throughout the nerve but did not co-localize with the βPS puncta or stripes (magenta, B').
Cross-sections of the peripheral nerve showing that Dlg5 does not co-localize with βPS

| Gene | Symbol | RNAi line | Glial phenotype |
|---|---|---|---|
| bazooka | baz | 38213 | Wild type |
| | | 35002 | Wild type |
| | | 39072 | Wild type |
| | | 2914 | Wild type |
| | | 2915 | Wild type |
| big bang | bbg | 36111 | Wild type |
| | | 15975 | Wild type |
| canoe | cno | 38194 | Wild type |
| | | 33367 | Wild type |
| CASK ortholog | CASK | 35309 | Wild type |
| | | 51721 | Wild type |
| | | 32857 | Wild type |
| connector enchancer of kar | cnk | 31759 | Wild type |
| | | 33366 | Wild type |
| drop out | dop | 44547 | Wild type |
| | | 36059 | Wild type |
| discs large 1 | dlg1 | 35286 | Wild type |
| | | 33620 | Abnormal inner glial membrane |
| | | 34854 | Abnormal glial membrane |
| | | 31520 | Inner glia aggregates |
| | | 25780 | Inner glia aggregates |
| discs large 5 | dlg5 | 41832 | Abnormal inner glia membrane |
| | | 101596 | Abnormal inner and outer glia membrane |
| | | 46234 | Abnormal inner and outer glial membrane |
| | | 22496 | Wild type |
| Dishevelled | dsh | 31306 | Wild type |
| | | 31307 | Abnormal glia |
| dyschronic | dysc | 110019 | Wild type |
| Efa6 pleckstrin and Sec7 domain containing | Efa6 | 42321 | Wild type |
| Fife | Fife | 110099 | Wild type |
| Glutamate receptor binding protein | Grip | 40930 | Wild type |
| | | 41978 | Wildtype |
| Grasp65 | Grasp65 | 34082 | Wild type |
| HtrA2 | HtrA2 | 28544 | Wild type |
| | | 55165 | Wild type |
| Inactivation no afterpotential d | InaD | 26211 | Wild type |
| inturned | in | 27252 | Wild type |
| Kermit | kermit | 53349 | Wild type |
| | | 109297 | Abnormal glial membrane |
| Lap-1 | Lap-1 | 27036 | Wild type |
| | | 18600 | Wild type |
| LIM-kinase1 | LIMK1 | 26294 | Abnormal glial membrane |
| | | 42576 | Wild type |
| locomotion defects | loco | 32456 | Wild type |
| | | 110275 | Abnormal glial membrane and swellings |
| | | 9248 | Abnormal glial membrane and swellings |
| menage a trois | metro | 35810 | Wild type |
| | | 110814 | Wild type |
| | | 29967 | Wild type |
| Myosin heavy chain like | mhcl | 51456 | Wild type |
| Magi | magi | 51761 | Wild type |
| | | 25792 | Wild type |
| | | 35279 | Wild type |
| | | 33411 | Wild type |
| par-6 | par-6 | 35000 | Wild type |
| | | 37478 | Wild type |
| | | 38361 | Vacuole like structures near inner glia nuclei, abnormal glial membrane |
| | | 39010 | Wild type |
| | | 108560 | Wild type |
| | | 19731 | Wild type |
| | | 26282 | Wild type |

| | | | |
|---|---|---|---|
| Patj | Patj | 35747 | Wild type |
| | | 38193 | Wild type |
| PDZ-GEF | PDZ-GEF | 28928 | Wild type |
| | | 31248 | Wild type |
| PICK1 | PICK1 | 31258 | Wild type |
| Prosap | Prosap | 40929 | Wild type |
| | | 40909 | Wild type |
| | | 27284 | Wild type |
| Protosome specific GEF | PsGEF | 44061 | Wild type |
| | | 33433 | Wild type |
| Ptpmeg | Ptpmeg | 39007 | Wild type |
| | | 43246 | Wild type |
| polychaetoid | pyd | 33386 | Wild type |
| | | 28920 | Wild type |
| | | 35225 | Wild type |
| Rim | Rim | 44541 | Wild type |
| | | 55741 | Wild type |
| | | 27300 | Wild type |
| RhoGAP19D | RhoGAP19D | 106241 | Wild type |
| | | 43955 | Wild type |
| RhoGAP100F | RhoGAP100F | 106241 | Wild type |
| | | 32946 | Wild type |
| RhoGEF2 | RhoGEF2 | 34643 | Interglial membrane swelling and axon defasciculation |
| | | 31239 | Interglial membrane swelling and axon defasciculation |
| Rhophilin | Rhp | 110377 | Wild type |
| | | 24111 | Wild type |
| | | 54474 | Wild type |
| scribbled | scrib | 35748 | Abnormal glial membrane |
| | | 39073 | Wild type |
| | | 29552 | Abnormal glial membrane |
| site-2 protease | S2P | 4601 | Wild type |
| | | 35760 | Glial swelling and axon defasiculation |
| Slip1 | Slip1 | 101106 | Wild type |
| | | 33007 | Inner glial aggregates |
| | | 51724 | Wild type |
| Spinophilin | Spn | 105888 | Wild type |
| | | 19658 | Wild type |
| sprite | sprt | 107873 | Wild type |
| SRY interacting protein 1 | Sip1 | 109289 | Wild type |
| | | 16958 | Wild type |
| short spindle 6 | ssp6 | 107399 | Wild type |
| | | 15622 | Wild type |
| still life | sif | 25789 | Wild type |
| | | 27406 | Wild type |
| stardust | sdt | 35291 | Wild type |
| | | 33909 | Some nerves with abnormal inner glial membrane |
| | | 33991 | Inner glia aggregates and abnormal outer glial membrane |
| | | 37510 | Abnormal inner and outer glial membrane |
| Syntrophin-like 1 | Syn-1 | 27893 | Wild type |
| | | 27504 | Wild type |
| Syntrophin-like 2 | Syn-2 | 28363 | Wild type |
| | | 42601 | Wild type |
| | | 31556 | Wild type |
| | | 34594 | Wild type |
| | | 34725 | Some pinched nerves |
| Varicose | vari | 28599 | lethal |
| | | 35193 | lethal |
| veli | veli | 29590 | Wild type |
| | | 110556 | Wild type |

| | | | |
|---|---|---|---|
| | | 46963 | Wild type |
| X11L | | 29309 | Wild type |
| Z-band alternatively spliced PDZ –motif protein-motif 52 | Zasp52 | 31561 | Wild type |
| | | 106177 | Wild type |
| Z-band alternatively spliced PDZ –motif protein-motif 66 | Zasp66 | 102980 | Wild type |
| Z-band alternatively spliced PDZ –motif protein 67 | Zasp67 | 103225 | Wild type |
| | | 42874 | Glial swelling and axon defasiculation |
| CG34375 | CG34375 | 101586 | Wild type |
| | | 23904 | Wild type |
| CG42312 | CG42312 | 101585 | Wild type |
| CG42788 | CG42788 | 108180 | Wild type |
| | | 100279 | Wild type |
| | | 22311 | Wild type |
| | | 31556 | Wild type |
| | | 34594 | Wild type |
| | | 34725 | Wild type |
| CG6688 | CG6688 | 106372 | Wild type |
| | | 48261 | Wild type |
| CG10362 | CG10362 | 8317 | Wild type |
| CG15617 | CG15617 | 19149 | Wild type |
| | | 104556 | Wild type |
| CG15803 | CG15803 | 51449 | Wild type |
| CG32758 | CG32758 | 38225 | Wild type |
| CG3402 | CG3402 | 110191 | Wild type |
| | | 21485 | Wild type |
| CG43707 | CG43707 | 102743 | Wild type |
| CG9588 | CG9588 | 28527 | Wild type |
| | | 100126 | Wild type |
| | | 47763 | Wild type |
| CG5921 | CG5921 | 37875 | Wild type |

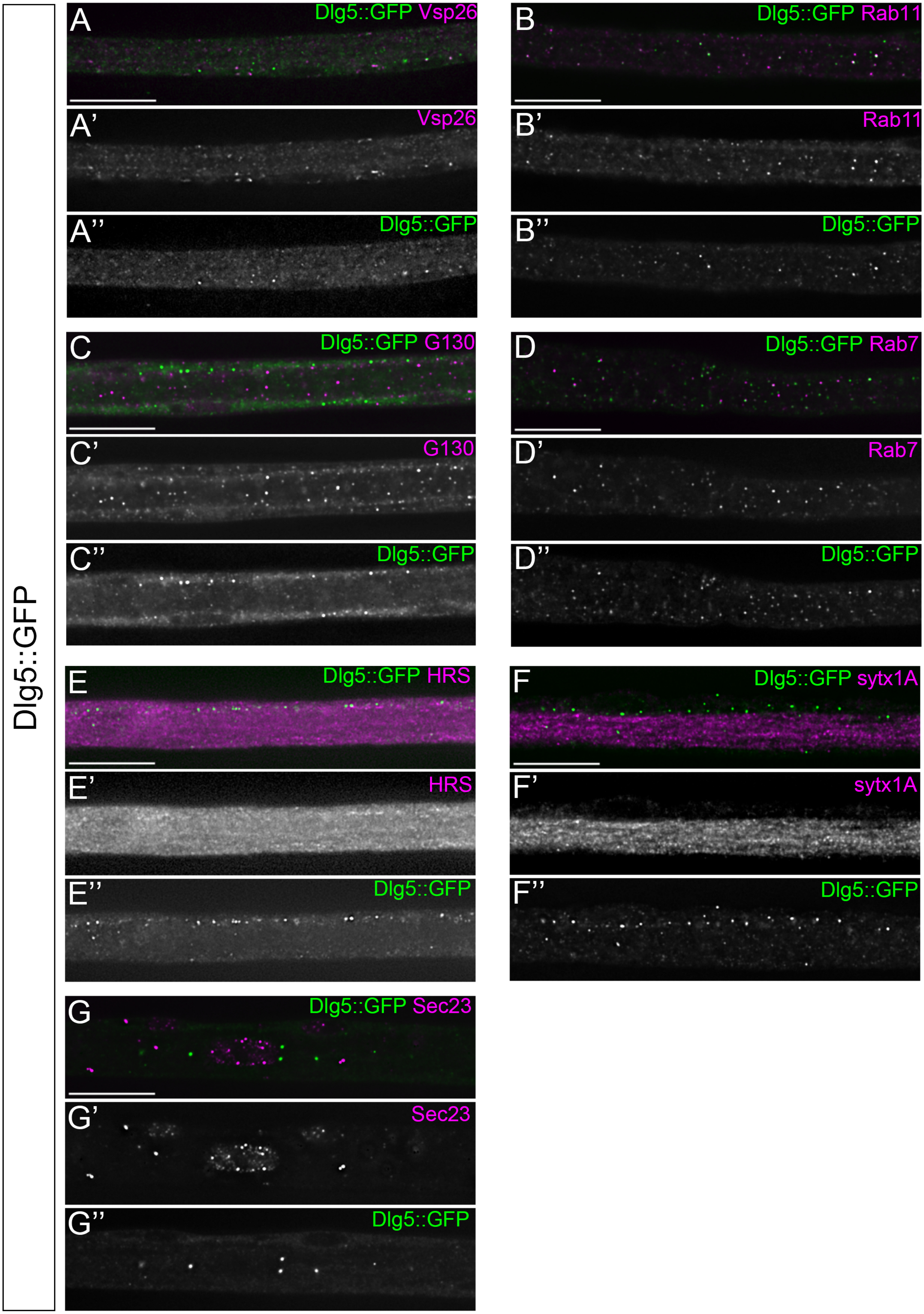
Dlg5::GFP
A
Dlg5::GFP Vsp26
A'
Vsp26
A''
Dlg5::GFP
B
Dlg5::GFP Rab11
B'
Rab11
B''
Dlg5::GFP
C
Dlg5::GFP G130
C'
G130
C''
Dlg5::GFP
D
Dlg5::GFP Rab7
D'
Rab7
D''
Dlg5::GFP
E
Dlg5::GFP HRS
E'
HRS
E''
Dlg5::GFP
F
Dlg5::GFP sytx1A
F'
sytx1A
F''
Dlg5::GFP
G
Dlg5::GFP Sec23
G'
Sec23
G''
Dlg5::GFP

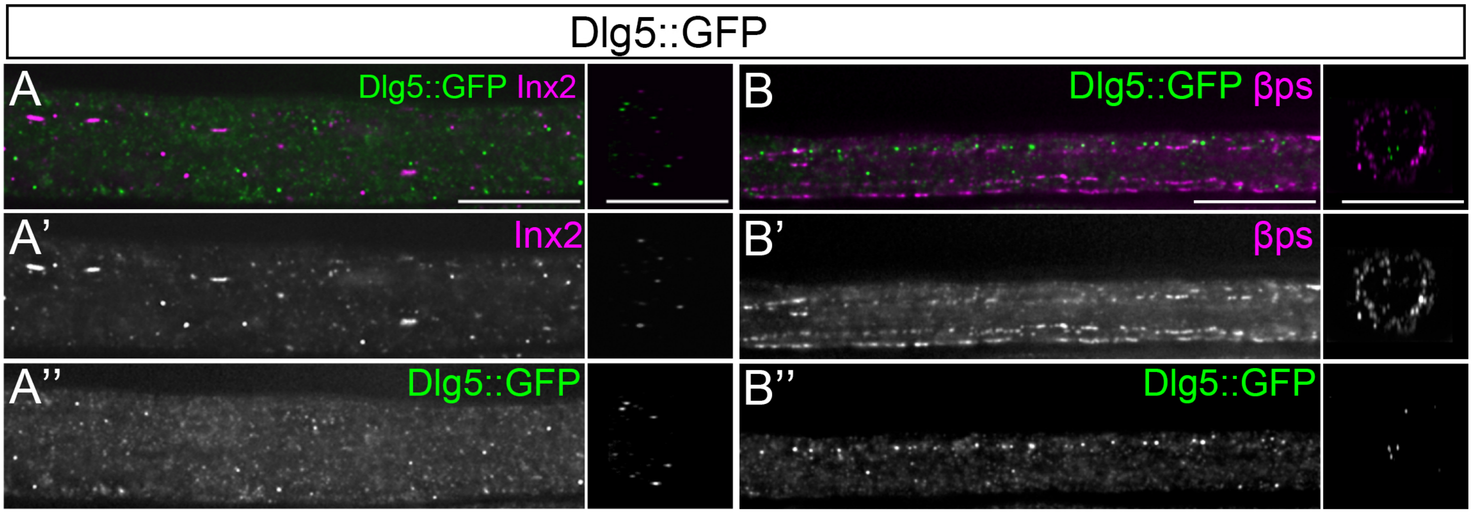
Dlg5::GFP
A
Dlg5::GFP Inx2
A'
Inx2
A''
Dlg5::GFP
B
Dlg5::GFP βps
B'
βps
B''
Dlg5::GFP
